# High residual pesticide contamination despite contrasted feeding treatments in semi-captive bred grey partridges

**DOI:** 10.64898/2026.07.23.740315

**Authors:** Sophie M. Dupont, Karine Monceau, Agathe Gaffard, Anaïs Rodrigues, Maurice Millet, Julie Noualhier, Audrey Bailly, Vincent Bretagnolle, Olivier Pays, Jérôme Moreau

## Abstract

Food security is one of the major ongoing global anthropic challenges, driving the intensification of agricultural systems notably through the widespread use of phyto-pharmaceutical products (PPPs). Although effective for maximizing production yields through pest control, PPP contamination is pervasive across the entire agroecosystem, raising concerns about human and non-target species exposure and health effects. Organic farming has therefore been promoted as a sustainable alternative to ensure human food security and mitigate these impacts. Yet, the pathways of PPP non-target organisms’ contamination are insufficiently understood. While most research has focused on ingestion as the primary route of exposure, increasing evidence suggests more complex contamination dynamics. In this study, we investigated the role of food intake in shaping PPP contamination profiles in semi-captive grey partridges (*Perdix perdix*). To do so, 80 birds were housed in open aviaries within an agricultural landscape and fed either conventional or organic grains over a five-month period. After drawing their PPP blood profiles at the end of the exposure period, we assessed their loads and compared the pesticide prevalence and abundance between food treatments. We found similar quantity and diversity of PPP residues in blood from conventionally- and organically-fed partridges, with few differences in specific PPP compounds. Overall, these findings suggest that food intake is not necessarily the only source of contamination for non-target organisms, with alternative pathways (e.g., inhalation, dermal contact, soil and water exposure) likely playing a substantial role. Integrating multi-pathway contamination monitoring across trophic levels within a One Health framework would now be essential to better understand and mitigate PPP contamination risks.

## 1. INTRODUCTION

Food security is one of the key global stakes for ensuring equitable feeding of the growing human population (which is expected to reach 9 to 10 billion people by 2050; Godfray et al., 2010). To do so, agricultural systems have evolved since the post-World War II period to maximize food production through higher yields *per* unit area. In this line, chemical inputs, and notably Phyto-Pharmaceutical Products (PPPs), have been used as part of conventional agricultural management strategies to control pests, i.e., fungi, insects, and weeds, that threaten food integrity. These PPPs are used as preventive or curative measures, meaning they are applied year-round, with timing varying according to seasonal agricultural constraints, alone or as part of mixtures. Concerningly, they are composed with an active chemical molecule accompanied by co-formulants stabilizers with physico-chemical properties inducing their possible mid-to long-term persistency in the environment (e.g., the insecticide Chlordecone and organochlorine pesticides, such as DDT, Dieldrin, and Heptachlor, exhibit very long environmental half-lives beyond two years, depending on the matrices and environmental conditions, Toan et al., 2009; Saaidi et al., 2023) but also their transferability throughout the biological compartments. Indeed, PPPs can be either bioaccumulated (i.e., stored within organisms in different tissues) and/or biomagnified (i.e., accumulated along the trophic chain) by organisms throughout ecosystemic transfers (Tison et al. 2024). Altogether, it leads to the contamination of the entire agroecosystem: air, soil, water, and organisms (Bonmatin et al. 2015; Socorro et al. 2016; Hladik et al. 2018; Giorio et al. 2021; Mauser et al., 2025).

Organic farming was then promoted to integrate a sustainable facet for human food security, especially necessary in our changing world. Concretely, new agricultural pest management practices, such as crop rotation, natural predators, and resistant plant varieties thereby banning chemical use (Stein-Bachinger et al. 2020), were developed and indirectly help to mitigate PPP–related environmental and non-target species contamination. While representing a significant potential lever to tackle the pervasive issue of PPPs contamination stemming from conventional agriculture, recent investigations have, however, revealed that non-target vertebrate species can exhibit PPP contamination even when inhabiting organic agricultural areas (e.g., Pelosi et al. 2021; Brodeur et al. 2022; Fritsch et al. 2022; Mauser et al., 2025). Indeed, some chemicals can travel up to 1,000 km from their application site through air and clouds, mainly due to spraying techniques and dust drift (Shen et al., 2005; Bianco et al., 2025), implying that pesticide-free crops may not necessarily exist (Mauser et al., 2025). This also highlights that there remain challenges in understanding the persistence of PPPs in the environment (air, soil, water, and organisms) and in elucidating the contamination pathways that predominate across the different compartments of ecosystems.

Non-target organisms living in agroecosystems may be contaminated by PPPs through various pathways (Botías et al. 2016; Wood & Goulson 2017). Most studies have focused on absorptive contamination by the internal digestive organs through food or water, as it is considered the primary mode of exposure to PPPs for vertebrates, (e.g. in birds: Mineau 2011; Lopez Antia et al. 2016; Moreau et al. 2022b). However, a growing number of studies are questioning the (key) role of the ingestion pathway in shaping the contamination profile and raised the urgent requirement of further exploration of the establishment of PPP contamination profiles as route of contamination seems more complicated (Faburé et al., 2025; Dupont et al. 2026). Indeed, organisms can also become contaminated by inhalation of chemical compounds when PPPs are applied to crops by spraying as well as by dermal (adsorptive) exposure, either through their skin, feathers, hair, or scales (Weltje et al., 2018; Moreau et al. 2022b; Faburé et al., 2025).

Because a pesticide-free ecosystem may not exist, one strategic approach to work on that contamination pathway question would be to compare the PPP contamination of non-target species consuming conventional (with pesticides) or organic (without pesticides) food in semi-controlled conditions, such as mesocosms or captive settings. This would allow specific pathways of PPP exposure through food ingestion to be manipulated and isolated, while keeping other parameters constant, i.e., species would still be exposed to water and air absorption, as well as adsorptive routes of contamination. It will then help us, once and for all, assess the importance of the ingestion pathway in organism impregnation and explore valuable insights for ecotoxicological mitigation. Here, we specifically tested whether providing conventional *vs* organic food influences PPP contamination profile in semi-captive grey partridges (*Perdix perdix*) housed in open aviaries located in an agricultural landscape, i.e., in close connection with the surrounding habitat (air, soil, and water runoff). This farmland bird species presents an adaptability to captivity, enhancing its suitability as a model organism in experimental setups. Moreover, studying the grey partridge is particularly relevant in an ecotoxicological context as high-input agriculture has been identified as one of the most influential pressures explaining its population declines (e.g., Chiron et al. 2014; Li et al. 2020; Rigal et al. 2023). In brief, previous experimental studies already highlighted sublethal effects on their physiology, notably a sex-based impact on body condition and behavior after ingestion of conventional grains containing PPP residues (Moreau et al. 2021; Gaffard et al. 2022; Dupont et al., 2026). Concretely, 80 immature semi-captive birds from a balanced sex-ratio were fed with conventional (3 PPPs quantified) or organically-treated (no PPP quantified) grains for five months. Following the feeding treatment, we conducted a minimally invasive, multiplex analysis of PPP residues in blood to quantify exposure to 94 PPPs across different pesticide families and draw individual contamination profiles, i.e., which pesticides each individual is contaminated by. We first predicted that, if ingestion uptake is the main contamination pathway, there would be a dietary “contamination print” on the individual profile of conventionally-fed partridges, associated with the PPPs quantified in food consumed, while organically-fed individuals would present little or no PPP residues. We then expected substantial differences in contamination profiles between our two feeding groups, as well as greater variability among conventionally-fed individuals compared to organically-fed ones, due to their broader potential sources of contamination. Accordingly, we hypothesized a high prevalence and abundance of certain pesticides, particularly those present in the provided grains, in conventionally-fed partridges and very low prevalence and abundance of all pesticides in organically-fed birds. On the other hand, if other absorptive and adsorptive sources of contamination from their surrounding agricultural environment are (at least) as important as food ingestion, we predicted similar PPP contamination profiles for our two feeding groups. In such case, we expected limited dissimilarity in individual contamination profiles between our two experimental groups and comparable inter-individual variability. We also hypothesized similar pesticide-specific prevalence and abundance between conventional and organic feeding treatments, with possibly a few exceptions directly associated with feed contamination, notably.

## 2. MATERIALS AND METHODS

All experiments complied with French legislation on animal experimentation, and all experimental protocols were approved by the Committee of Animal Experimentations of the Deux-Sèvres French District (APAFIS#9465-201703101551625).

### 2.1. Experimental design

The five-month experimental protocol was conducted from Autumn 2020 to Spring 2021 at a commercial game farm in an agricultural area in south-western France (Deux-Sèvres, France; Dupont et al., 2026). In detail, the two adjoining pens (each measuring 100 m × 10 m × 4 m) were located on a plot of land largely surrounded, in decreasing order of area coverage, by cereal crops, meadows/pastures, sunflowers, fodders, oleaginous crops, fallow lands, and hedgerows (Figure 1). These agricultural lands together covered 86.79% of the environment within a 1-km radius in 2020, with the 13.21% remainder corresponding to forests, urbanized areas, and roads. Among the neighboring agricultural parcels, 99.02% were reportedly managed under conventional practices, meaning that only 0.98% were declared organic to the Common Agricultural Policy - Ministry of Agriculture (Figure 1).

**Figure 1.**
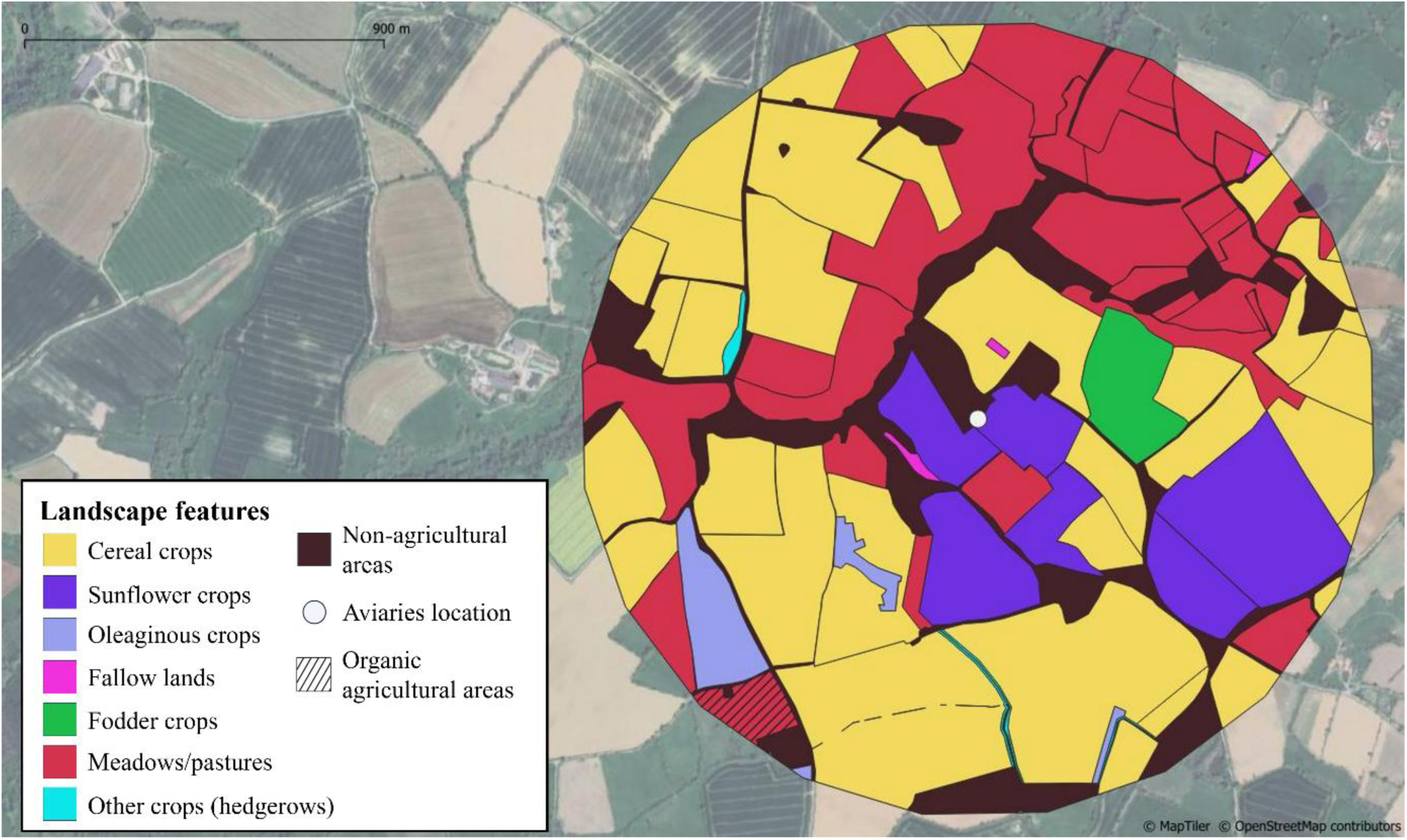
Commercial game farm location where the food experiment took place from Autumn 2020 to Spring 2021 with 80 seven-month-old grey partridges. Landscape features within a 1-km radius of the aviaries are derived from the government’s Graphical Parcel Register for 2020. The hatched area corresponds to plot registered as farmed using organic farming methods. Dark areas represent non-agricultural areas, i.e., forests, urbanized areas, and connecting zones.

The 80 seven-month-old grey partridges (40 females and 40 males, sexed based on their unambiguous visual secondary sexual characters appearing after their first molt) monitored in this study were randomly selected among a common garden group of 200 unrelated partridges, corresponding to the F3 generation of wild trapped birds. Importantly, before the start of the experimental protocol, they all lived in the same pen at the commercial game farm and were fed an adequate commercial juvenile poultry feed from birth (STARGIB Entretien; see Appendix 1 for its composition, and Figure 2). The 40 individuals from the conventional group are those described in Dupont et al. 2026, in which the link between contamination patterns and health was examined.

When the experiment started, the 80 partridges were individually identified with an alphanumeric metal ring and then randomly assigned in pairs (balanced sex ratio) to one of our two experimental treatments: conventional or organic food. Each group was represented by a housing pen equipped with feeders, drinkers, and metal sheets with a wooden structure for shelters. No dogs or cats were allowed inside the aviaries, but it is impossible to be certain that domestic, feral or wild animals did not come near the fence, as the area surrounding the pens was not enclosed. Importantly, the two side-by-side pens were built with mesh, separated only by a wire fence, and had muddy, grassy soil continuous with the surrounding area. In other words, all partridges were exposed to a natural light cycle and subject to the same abiotic environmental conditions across their entire habitat (air, soil, and water runoff), which were continuous with the neighboring agricultural environment. Moreover, identical sink water was daily provided to the two experimental groups *ad libitum*. To note, no PPP contamination profile was drawn for air, aviary soil, or sink water. Birds received the equivalent mix of seeds (wheat, corn, pea, and faba beans in equal quantities) obtained from either conventional (with pesticides, hereafter defined as our ‘conventional’ group) or organic (without pesticides, hereafter designated as our ‘organic’ group) agricultural practices. In detail, grains from the conventional group were purchased from producers applying various PPPs during the cropping season (see Appendix 2 for the list of PPPs applied by the farmers to crops); while those from the organic treatment were harvested from certified organic crops. Our experimental protocol was validated by a multi-residue gas chromatography mass spectrometry analysis, drawing the list of chemical substances quantified in conventional and organic grains (see protocol described below and Table 1 for the list of chemicals detected in food provided and associated quantity). During winter, 11 partridges inadvertently died (five conventionally- and six organically-fed birds respectively, representing 13.75% of global mortality), which is common in breeding farms (J. Blandin, personal communication).

### 2.2. Data collection

In March 2021, i.e., after the five-month food treatment (i.e., conventional *vs* organic), morphometric measurements were taken on the 69 remaining birds (N_conventional_ = 35, 18 females and 17 males and N_organic_ = 34, 18 females and 16 males). In detail, we measured the lengths of the left and right tarsus using a caliper (accuracy: ± 0.10 mm) and recorded body mass with a spring scale Pesola 500g (accuracy: ± 1.00 g).

A blood collection (50 µL) was also performed for pesticide analysis in the brachial vein using a sterile needle (Ø 0.06 mm) and heparinized micro-capillary tubes. Blood was immediately stored in Eppendorf tubes at -80°C until subsequent laboratory analyses (± three months later). Pesticide exploration has been conducted in this biological matrix, as it does not require sacrificing animals and will provide a short-term picture of PPP contamination (Espín et al., 2016). Moreover, in accordance with animal welfare principles, we minimized as possible stressful procedures by restricting the combination of capture, handling and blood collection to a single occasion at the very end of the experimental period. Here, all birds were born, reared under identical conditions, and received the same conventional food during wintering. The list of quantified chemical substances they were exposed to by this commercial juvenile poultry food has been drawn (see protocol described below) and is provided in Table 1. Consequently, we would consider that any noticeable difference in pesticide contamination between our two experimental groups to be attributable to the differential food provided during the five-month food treatment, particularly for those quantified in provided grains (conventional *vs* organic, Table 1).

### 2.3. Contamination profile

#### 2.3.1. Analytics for PPP residues in ingested seeds

Pesticide quantification of commercial juvenile poultry food and seeds from the two food treatments was conducted at the GIRPA laboratory (Angers, France) in October 2020, i.e., prior to the start of the experiment, to validate the experimental design of this study and establish the expected “contamination print” from our first prediction (see Results). A total of 585 molecules - including insecticides, fungicides, herbicides, and synergists - were searched for in pellet and grain samples. Among them, most PPPs assessed in blood partridges (see the detection/quantification protocol associated below) were included, except for 2,4-MCPA, Dimethenamid-P, Mecoprop-P which were searched in blood partridges but not in seeds. The contamination investigation was carried out independently for each seed composing the partridge experimental food, i.e., wheat, corn, pea, and faba bean, but the protocol was identical for all of them.

The methodology used for quantifying PPPs was developed based on QuEChERS NF 15662 (CECF, 2008). The process employed multiple reaction monitoring (MRM) mode, with dual mass transitions designated for each PPP compound, i.e., one for quantification and another for qualification. In detail, 2.00 g of blended pellet or grain was mechanically agitated with 10 mL acetonitrile, 10 mL ultrapure water, and 50 µL Chlorpyrifos-(diethyl-d10) [2 mg/L], i.e., an extraction tracer, for 20 minutes prior to transfer to a 50 mL QuEChERS tube containing the appropriate mixture of salts (magnesium sulfate, sodium chloride and citrate buffer salts). The tubes were then shaken vigorously for 1 min and centrifuged to separate the two phases, i.e., the acetonitrile and aqueous phases. From the acetonitrile phase, a Liquid Chromatography coupled with tandem Mass Spectrometry (LC-MS/MS, API 5500, Sciex, Toronto, Canada; with High-Performance Liquid Chain (HPLC) Shimadzu LC40AD and Synergy Hydro Reverse Phase (RP) column, Kyoto, Japan) analysis was first employed to detect and quantify acid chemicals. To do so, 10 µL formic acid, 10 µL diuron d6 and 230 µL ultrapure water were added to 250 µL supernatant. Then, quantification of PPPs from the acetonitrile phase involved an extract purification with sample transfer in a QuEChERS tube containing 150 mg PSA and 400 mg C_18_, but no Graphite carbon black (GCB), followed by a rotary evaporation step with 40 µL formic acid and 50 µL ethylene glycol in a Büchi tube before elution in 950 µL of ethyl acetate. Gas Chromatography coupled with tandem Mass Spectrometry (GC-MS/MS, GC7000 Agilent, Waldbronn, Germany; with High-Efficiency Source (HES), HP-5ms Ui 2×15m column and Multi-Mode Injectors (MMI)) analysis was then conducted to search for nonpolar chemicals. There, a 500 µL aliquot of extract was mixed with 10 µL alachlor-d13 (5 ppm). Finally, the remaining 250 µL extract solution were evaporated to dryness in nitrogen and diluted in 250 µL acetonitrile, 10 µL diuron-d6 and 240 µL ultrapure water before conducting a second LC-ESI-MS/MS analysis to search for purified chemicals.

The concentrations of each PPP in the sampled juvenile and conventional foods were quantified in milligrams *per* kilogram (mg.kg^-1^) and are presented in Table 1. No PPP were found in the Organic seeds provided to partridges. For all the compounds targeted, the limit of quantification (LOQ) was 0.01 mg.kg^-1^. When a PPP was not quantified in the food (<LOQ), the concentration was set to 0.00 mg/kg.

#### 2.3.2. Analytics for PPP residues in blood

The blood extraction and analysis processes involved several meticulous steps to ensure accurate quantification of pesticide residues, following the validated method reported by Rodrigues et al. (2023). For both LC-MS/MS and GC-MS/MS analyses, the MRM mode was used. Data acquisition and processing were performed using Excalibur software, and compound identification was based on retention times, ion transitions, and fragmentation patterns, as detailed in Appendix A.1 and A.2 from Rodrigues et al. (2023). In brief, blood samples were thawed, weighed, and spiked with 10 µL of carbendazim-d6 internal standard before homogenization. A liquid-liquid extraction protocol adapted from Goutner et al. (2011) was used, employing dichloromethane and ethyl acetate (1:1), followed by sonication and repeated three times. The supernatants were pooled, evaporated to a final volume of 500 µL, and stored at -20°C for dual analysis by GC-MS/MS and LC-MS/MS. A total of 94 molecules - including insecticides, fungicides, and herbicides - were searched for in all 69 blood samples. The complete list of the monitored compounds, along with validation parameters, is presented in Appendix 3.

For non-volatile compounds, samples were analyzed using LC-MS/MS equipped with a C_18_ column (Macherey-Nagel Nucleodur C_18_ Pyramid, 150 mm length × 3 mm inner diameter, 3 μm particle size). Separation was performed using a gradient of water and acetonitrile, both containing 0.1% formic acid, and detection was carried out in positive ion mode.

For volatile and semi-volatile compounds, extracts were deposited onto Tenax®-TA tubes and analyzed using a GC-MS/MS. A derivatization step with N-(tert-butyldimethylsilyl)-N-methyltrifluoroacetamide (MTBSTFA) as reagent was employed to improve the detectability of some semi-volatile compounds. Prior to separation, thermal desorption (TD) was performed using an ATD 350 system connected to the GC-MS/MS, with compound separation achieved on a capillary column (Macherey-Nagel OPTIMA XLB, 30 m length × 0.25 mm inner diameter, 0.25 µm film thickness) under a programmed temperature gradient.

The concentrations of each PPP detected in partridge blood samples were expressed in milligrams *per* kilogram (mg.kg^-1^) and are summarized in Table 2. For compounds detected at a level below the LOQ (see Appendix 3 for compound-specific threshold) but above the limit of detection (LOD, Appendix 3), a substituted value corresponding to the PPP-specific LOQ / 2 was assigned. When a PPP was not detected (< LOD), the concentration was set to 0 mg.kg^-1^. Such data handling represents a standard approach in contaminant analyses (see for example, Fritsch et al., 2022; Rico et al., 2021; Shin et al., 2020).

#### 2.3.3. Definition of PPP load

To compare the PPP load of partridges between our two experimental groups (conventional *vs* organic), we first determined the number of PPPs (N_PPP_) detected (> LOD) in the blood of each bird. In brief, while not accounting for the specific toxicity and quantity of each compound, this metric has been widely used in ecotoxicological studies as a useful proxy of individual cumulative risk, assuming that a higher number of PPPs increases the likelihood of additive and/or synergistic interactions between chemicals, and thus potential toxic risk (Zaller et al., 2022). Moreover, we calculated the total sum of scaled pesticide concentrations (Σ[pesticides]_scaled_) for each partridge, considering all pesticides quantified in at least one bird in the context of our study (N = 33, see Table 2 for the detailed list). Such approach provided us with a comparable estimate of the short-term cumulative pesticide content of all individuals of interest in the context of our study. It then reflected the overall individual level of impregnation as part of our sample, assuming that higher (positive) levels correspond to a possible greater toxic risk (Tartu et al., 2014; Fritsch et al., 2022; but see Hernández et al., 2017). Concretely, we first applied a centered-scaled transformation to the 33 detected substances and summed all the scaled concentrations measured *per* individual, without considering their respective toxicity level. Finally, we estimated the total toxicity index (TI_tot_) for each individual by calculating the sum of the ratio of each detected PPP concentration to its corresponding LD50 value. To note, for PPPs that were not detected in a given individual, the TI_PPP_ was equal to 0 as we used here the non-scaled PPP concentration in this calculation. This is currently the most widely used method in ecotoxicological studies to consider molecules’ toxicity (Dupont et al., 2026). Indeed, although this approach presents several methodological biases that are currently not issued (e.g., not considering potential synergism or antagonism in PPP mixtures, or including LD50 values originating from different species and corresponding to ingested PPP concentrations rather than blood-circulating ones), it still assesses the potential toxicity of the mixture an individual is impregnated by, with higher values being associated with a greater toxic risk of the mixture (Kortenkamp et al., 2009). The LD50 values were extracted from the Pesticide Properties DataBase. If the provided LD50 value exceeded a given value, e.g., 2000.0 mg.kg^−1^ (i.e., reported as ‘> 2000’), we considered the indicated value (on the example here: 2000.0 mg.kg^−1^) as the LD50 value in the calculation (see Appendix 3). We tested the relationship between our three PPP loads of interest using three distinct Pearson tests.

### 2.4. Statistical analysis

All statistical analyses were performed with RStudio software (version 4.3.0; R Core Team 2022). For all tests, *p*-values are given at the 5% significance level.

#### Body condition calculation

Because of the potential link between body condition and pesticides (see Moreau et al. 2021, for example), we included this morphometric variable as a covariate in all models testing the relationship between PPP loads and food treatment (see below). We assessed body condition by sex due to significant sexual dimorphism in this species. Average tarsus length was positively correlated with body mass (Pearson test, r_female_ = 0.460, *p-value*_female_ = 0.005, IC95% = [0.150; 0.628] and r_male_ = 0.640, *p-value*_male_ < 0.001, IC95% = [0.380; 0.806]). We then calculated the scaled mass index (SMI), implemented with the *smatr* package, using Peig & Green (2009) formula: SMI = 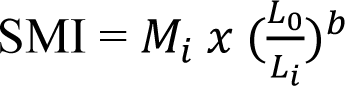, with M_i_ and L_i_ the body mass and the average tarsus length of the individual I respectively; L_0_ the arithmetic mean value of tarsus length for all females and males respectively (L_0,female_ = 43.07 mm, n = 36 females, L_0,male_ = 44.63 mm, n = 33 males); the exponent *b* the slope estimate of a standardized major axis regression of log-transformed body mass on log-transformed average tarsus length (b_female_ = 2.86, b_male_ = 1.97).

#### Comparison of prevalence and abundance for each PPP detected

First, for each PPP detected (> LOD) in a minimum of five individuals from one food treatment and in at least one from the other (to meet the minimum requirement for statistical test), we compared the number of detections and concentrations between food treatments (all results are presented in Table 2). First, samples detected levels (> LOD) of a given PPP were attributed a 1 and others a 0 and the number of detections were compared between food treatments (conventional *vs* organic) with either a Fisher exact test (if there were less than five positive samples in one food treatment) or a Chi-squared test (if there were more than five positive samples in both food treatments). Second, same samples (> LOD) of PPP were used to assess differences in the mean concentration of each quantified PPP between partridge food treatments (conventional *vs* organic) using Wilcoxon rank sum test.

#### Relationship between PPP loads and food treatment

We built three linear models to estimate the relationship between PPP loads (N_PPP_, Σ[pesticides]_scaled_ and TI_tot_) and the experimental food treatment (conventional *vs* organic). Each model included one PPP load feature as the variable of interest. We included food treatment (conventional *vs* organic), sex (female *vs* male), body condition and the ‘sex x body condition’ interaction as non-colinear explanatory fixed variables (VIF < 3). Following the statistical approach described in Dupont et al. (2026), we used AICc model selection based on ΔAICc < 2 as selection criteria and then full-model averaging estimate using the *MuMIn* package in the R software (Burnham et al., 2011). Model assumptions were visually checked.

#### Comparison of pesticide prevalence and abundance between food treatments

To compare contamination profiles between conventionally- and organically-fed individuals, we assessed differences in individual pesticide prevalence and abundance by calculating two complementary semi-parametric dissimilarity indices, i.e., the Jaccard and Bray-Curtis distances (Jaccard, 1912; Bray & Curtis, 1957). Concretely, the Jaccard index uses the presence or absence (1/0) of each compound, while the Bray-Curtis distance considers their concentration to determine complementary compositional differences in PPP contamination profiles, using the *vegdist* function from the vegan package (Oksanen et al., 2017). We then performed a permutational multivariate analysis of variance (PERMANOVA, *adonis2* function, vegan package) to test the overall effect of feeding treatments (conventional *vs* organic) on PPP profiles, with 1,000 permutations partitioning the sum of squared deviations between group centroids. We evaluated the effects of food treatments on dissimilarity patterns using the *envfit* function from the vegan package, with significance tested *via* a goodness-of-fit procedure based on 1,000 permutations. We further assessed variability in prevalence and abundance within each feeding group by testing homogeneity of dispersion using the *betadisper* function in the vegan package. We estimated whether group (conventional *vs* organic) dispersion relative to the centroids differed between our two food treatments using an ANOVA test with 1,000 iterations of free pairwise permutations tests.

## 3. RESULTS

### (a) Contamination pattern in foods and partridge blood

Commercial juvenile poultry food was contaminated by three out of the 585 molecules screened (i.e., Cypermethrin, Piperonyl butoxide and Pirimiphos-methyl, Table 1). In conventional seeds provided during the experiment, four PPPs were found (i.e., Chlorpyriphos-methyl, Piperonyl butoxide, Pirimiphos-methyl, and Tebuconazole). To note, commercial juvenile poultry food, and seeds were contaminated by two common molecules (i.e., Piperonyl butoxide and Pirimiphos-methyl). In organic food, no PPPs were quantified (Table 1).

**Table 1:**
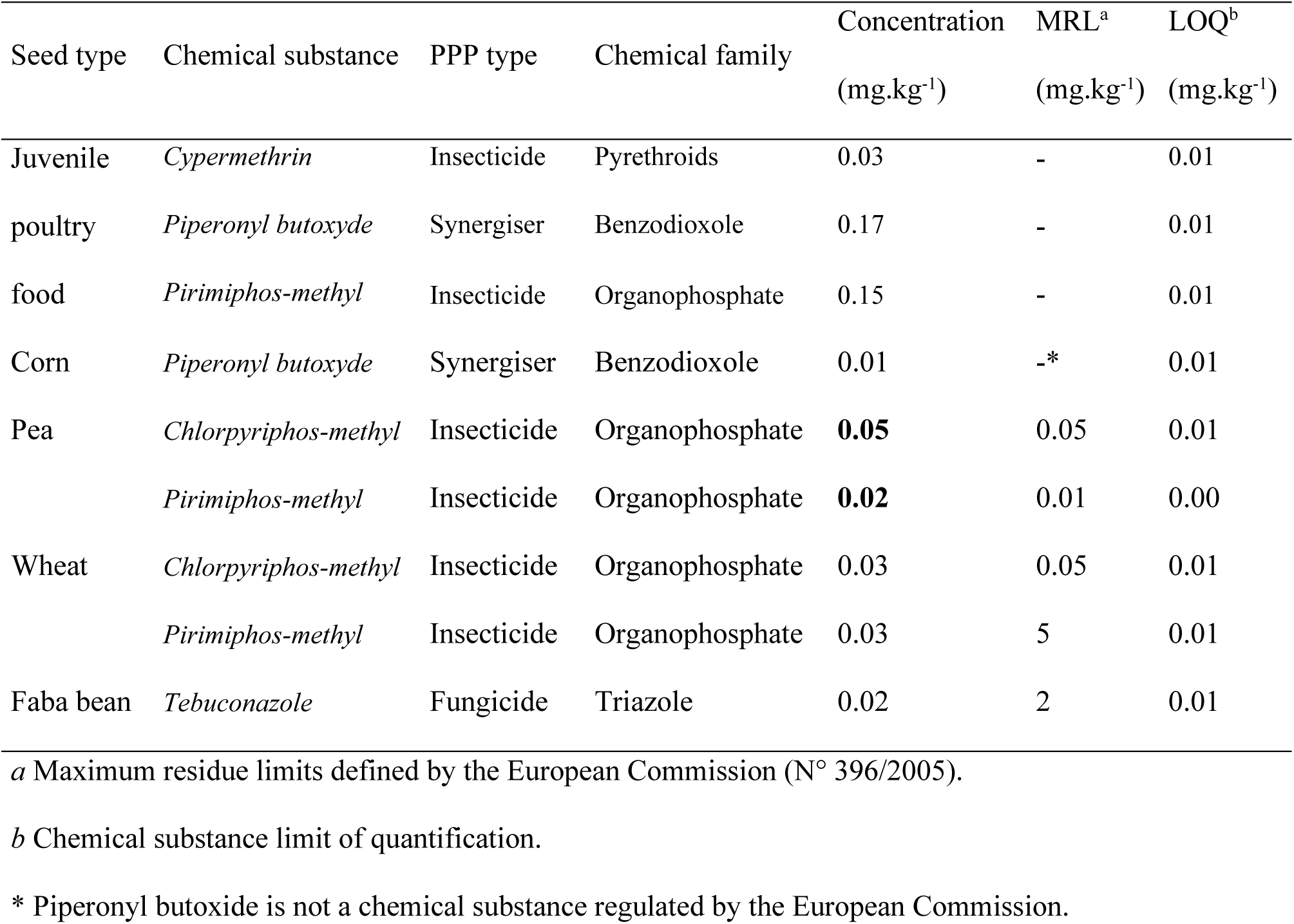
List of chemical substances quantified in both commercial juvenile poultry food and corn, pea, wheat and faba bean provided to conventionally-fed partridges using a multi-residue gas chromatography mass spectrometry (GC-MS) analysis (ISO 17025) carried out by the GIRPA laboratory (Beaucouzé, Loire Atlantique, France). Values in bold correspond to levels exceeding the current MRLs^a^.

| Seed type | Chemical substance | PPP type | Chemical family | Concentration<br>(mg.kg<sup>-1</sup>) | MRL<sup>a</sup><br>(mg.kg<sup>-1</sup>) | LOQ<sup>b</sup><br>(mg.kg<sup>-1</sup>) |
| --- | --- | --- | --- | --- | --- | --- |
| Juvenile<br>poultry<br>food | *Cypermethrin* | Insecticide | Pyrethroids | 0.03 | - | 0.01 |
|  | *Piperonyl butoxyde* | Synergiser | Benzodioxole | 0.17 | - | 0.01 |
|  | *Pirimiphos-methyl* | Insecticide | Organophosphate | 0.15 | - | 0.01 |
| Corn | *Piperonyl butoxyde* | Synergiser | Benzodioxole | 0.01 | -* | 0.01 |
| Pea | *Chlorpyrifos-methyl* | Insecticide | Organophosphate | **0.05** | 0.05 | 0.01 |
|  | *Pirimiphos-methyl* | Insecticide | Organophosphate | **0.02** | 0.01 | 0.00 |
| Wheat | *Chlorpyrifos-methyl* | Insecticide | Organophosphate | 0.03 | 0.05 | 0.01 |
|  | *Pirimiphos-methyl* | Insecticide | Organophosphate | 0.03 | 5 | 0.01 |
| Faba bean | *Tebuconazole* | Fungicide | Triazole | 0.02 | 2 | 0.01 |

<sup>a</sup> Maximum residue limits defined by the European Commission (N° 396/2005).
<sup>b</sup> Chemical substance limit of quantification.
\* Piperonyl butoxide is not a chemical substance regulated by the European Commission.

Among all partridges (conventional and organic), 33 out of the 94 PPPs screened in partridge blood were detected in at least one blood sample, including 15 fungicides, 13 herbicides and 5 insecticides (Figure 2; Table 2). Four PPPs were found in the blood of more than half of the individuals (Figure 2; Table 2): Bifenox, Diphenylamine, Nitenpyram and Tolylfluanid were detected in 95.6%, 81.2%, 71.0% and 56.5% of individuals, respectively.

Partridge contamination profiles were poorly congruent with both commercial juvenile poultry food, and seeds ones (Figure 2). Tebuconazole was detected once in each experimental group and Piperonyl butoxide twice in the organic group (Table 2). To note, Tebuconazole and Piperonyl butoxide levels reported in partridge blood were higher than those measured in foods (i.e., Tebuconazole = Food_conventional seeds_: 0.17 mg.kg^-1^, Blood_conventionally-fed partridges_: 1.32 mg.kg^-1^ and Blood_organically-fed partridges_: 1.42 mg.kg^-1^ ; Piperonyl butoxide = Food_juvenile_: 0.17 mg.kg^-1^, Food_conventional seeds_: 0.01 mg.kg^-1^, Blood_organically-fed partridges_: 8.53 ± 1.53 mg.kg^-1^). However, Chlorpyriphos-methyl, Cypermethrin, and Pirimiphos-methyl were not detected in any partridge.

**Figure 2:**
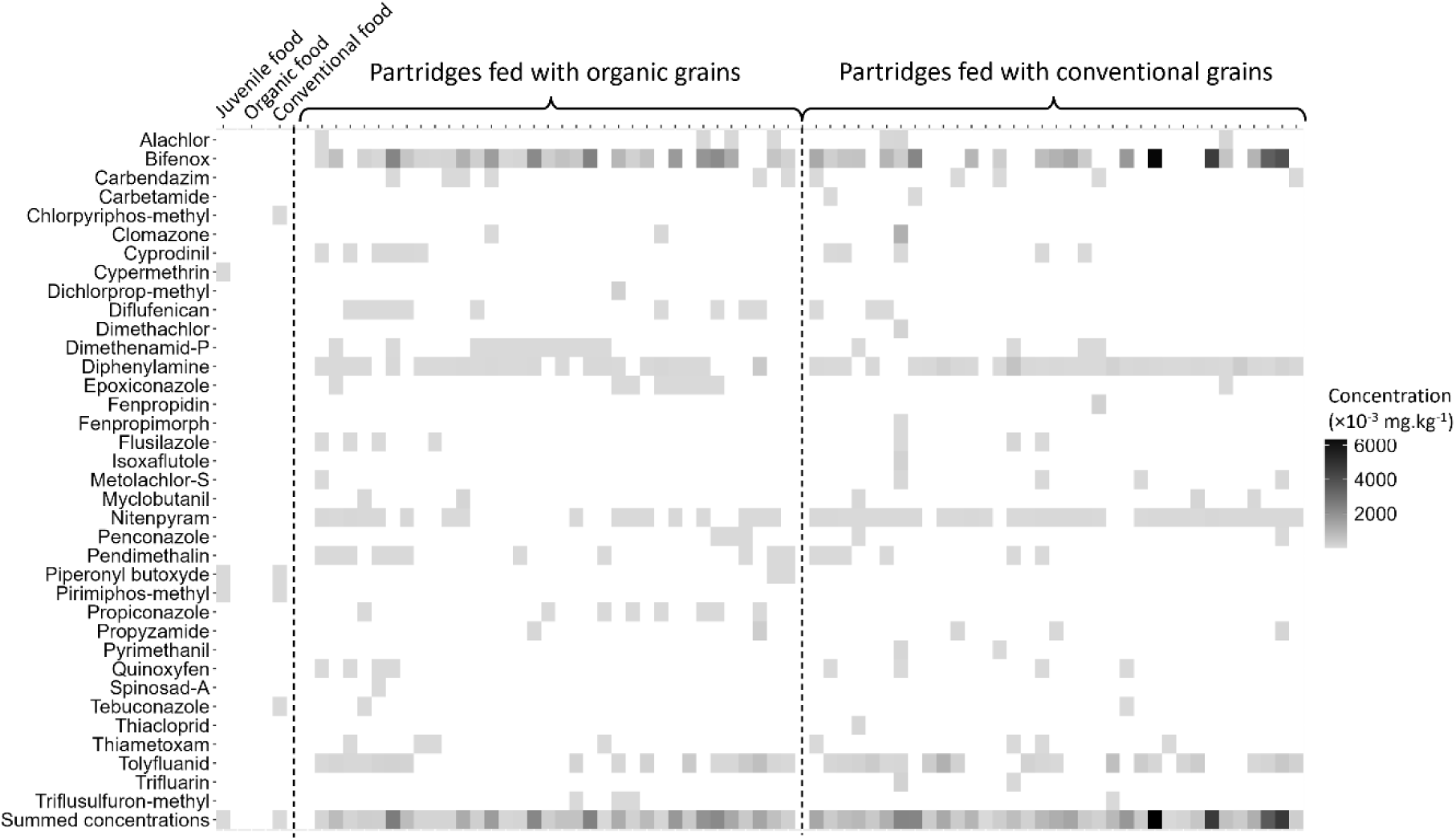
Heatmap of the distribution of phyto-pharmaceutical products’ (PPPs) concentration in food provided (juvenile pellets and experimental mix of seeds without or with pesticides in organic and conventional treatments respectively) and in partridge blood, organized according to food treatment (organic *vs* conventional). Here, we present the distribution of the 33 quantified out of 94 searched-for pesticides in the blood of 69 partridges in alphabetic order, alongside the list of PPPs found in food. PPP concentrations ranged from 0.01 to 6382.18 ×10^-3^ mg.kg^-1^. A value of zero was attributed when the PPP concentration was < LOD.

**Table 2:**
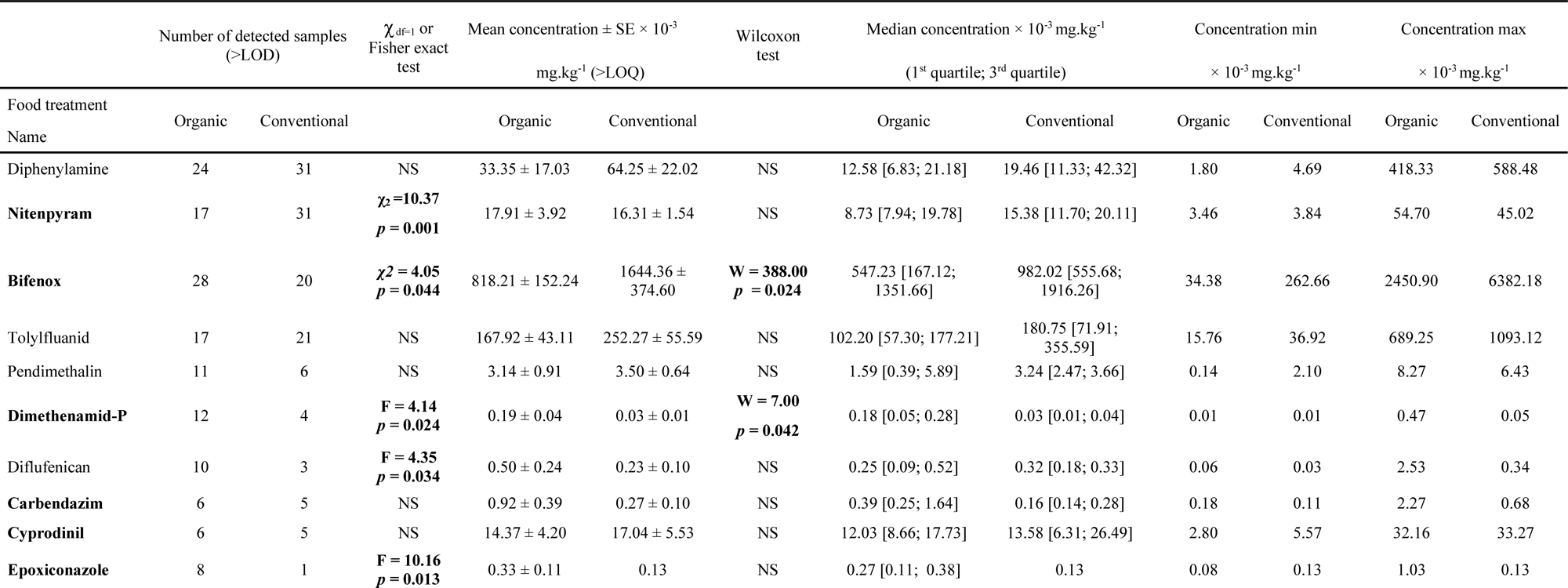
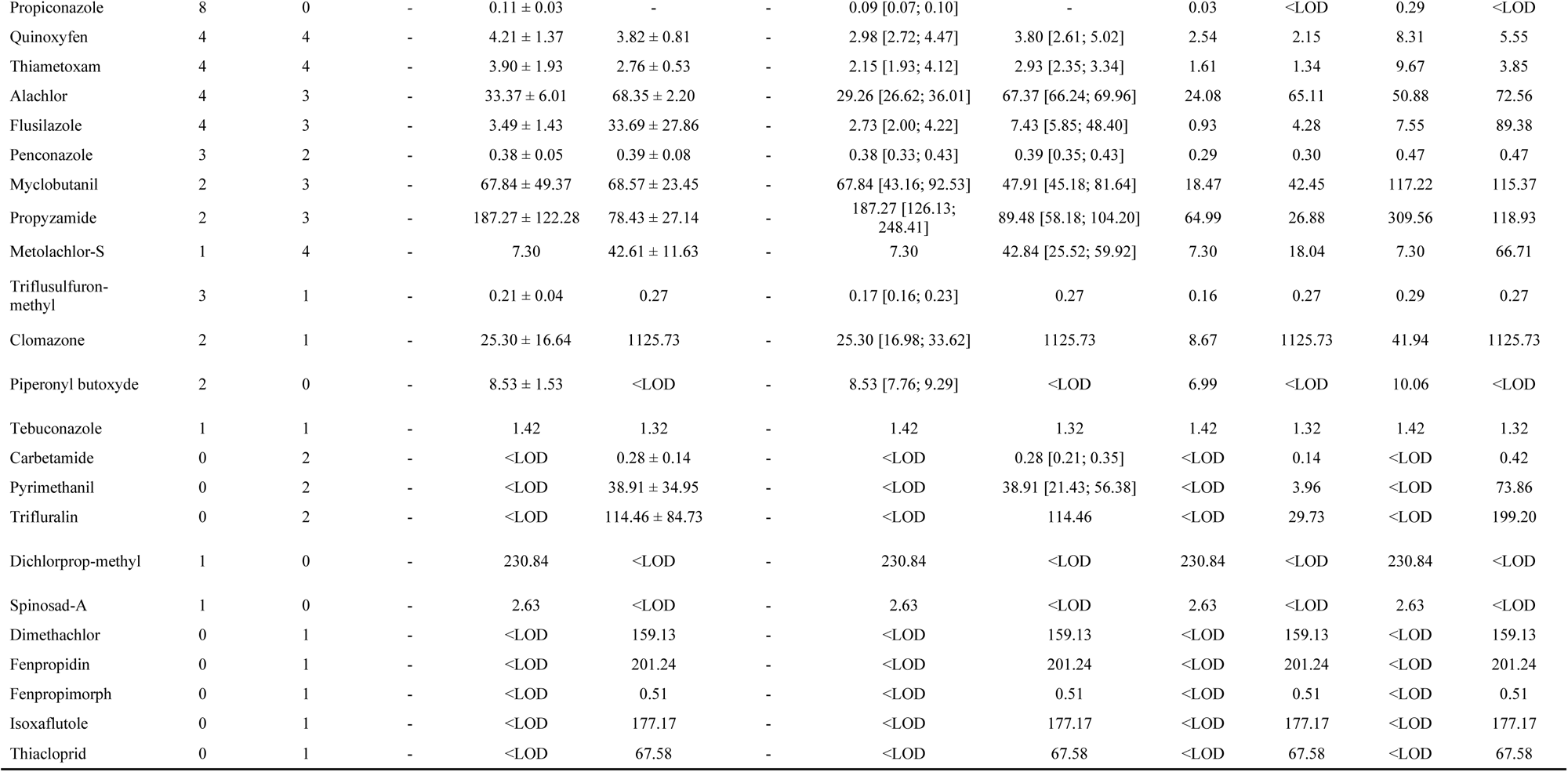
Number of detected samples, quantified mean concentration (± standard error, SE), median (1^st^ quartile; 3^rd^ quartile), minimum, and maximum for the 33 PPPs detected in 69 partridge blood samples according to food treatment (i.e., conventional, N = 35, or organic, N = 34). For each PPP detected (> LOD) in a minimum of five individuals from one food treatment and in at least one from the other, Chi-squared tests (when the number of detected samples > 5 for both food treatment) or Fisher exact tests (when the number of detected samples < 5 in one food treatment) were used. Wilcoxon tests were used for all PPPs where the number of quantified samples was > 5 in one food treatment and at least 1 on the other. Significant differences in prevalence or abundance respectively are in bold.

Overall, across both food groups combined, the N_ppp_ *per* individual ranged from 1 to 14 detected molecules (Table 3). Partridge Σ[pesticides]_scaled_ ranged from -8.30 to 61.80 mg.kg^-1^ (Table 3). Partridge TI_tot_ values ranged from 5.96×10^-6^ to 3.26×10^-2^ (Table 3).

N_PPP_ was positively correlated with Σ[pesticides]_scaled_ (Pearson test, r_Pearson_ = 0.794; CI95%: [0.686, 0.867], *p* < 0.001), but no correlation was found with TI_tot_ (r_Pearson_ = 0.095; CI95%: [-0.145, 0.324], *p* = 0.439). Σ[pesticides]_scaled_ was positively correlated with TI_tot_ (r_Pearson_ = 0.292; CI95%: [0.060, 0.495], *p* = 0.015).

**Table 3:** Descriptive information for the three PPP load features (N_PPP_, Σ[pesticides]_scaled_, TI_tot_) included as explanatory variables in the models. We presented here Minimum-Maximum and Mean ± standard error for all birds, i.e., conventionally- and organically-fed partridges respectively.

| | $N_{\text{PPP}}$ | | |
| --- | --- | --- | --- |
|  | All birds | Conventional | Organic |
| Min -Max | 1.00 - 14.00 | 1.00 - 14.00 | 3.00 - 10.00 |
| Mean ± SE | $5.30 \pm 0.26$ | $5.14 \pm 0.40$ | $5.47 \pm 0.33$ |
| | $\Sigma[\text{pesticide}]_{\text{scaled}}$ | | |
|  | All birds | Conventional | Organic |
| Min -Max | -8.30 - 61.80 | -8.30 - 61.80 | -8.05 - 14.83 |
| Mean ± SE | $0.00 \pm 1.14$ | $0.17 \pm 2.01$ | $-0.17 \pm 1.08$ |
| | $\text{TI}_{\text{tot}} \times 10^{-3}$ | | |
|  | All birds | Conventional | Organic |
| Min -Max | 0.01 - 3.26 | 0.01 - 3.26 | 0.02 - 1.35 |
| Mean ± SE | $0.56 \pm 0.08$ | $0.70 \pm 0.14$ | $0.42 \pm 0.07$ |

Based on the AICc model selection and full-model averaging estimate, no significant difference in the N_ppp_, nor the Σ[pesticides]_scaled_, nor the TI_tot_ were found between conventionally-and organically-fed partridges (Figure 3ABC, Table 4). Moreover, no sex or body condition effects on PPP loads were observed (Table 4).

**Table 4:** Grey partridge PPP load according to experimental food treatment (conventional *vs* organic), sex (female *vs* male), body condition and the interaction ‘sex x body condition’. Tests were performed using linear models and obtained from a model-averaging procedure. Significant effects are in bold type. Z-value and p-value are reported for all statistical tests. Parameter estimates, and standard errors, are provided here with the 95% confident interval. For TI_tot_ only, these four values are expressed as x 10^-3^.

|  |  | Estimate | SE | Adjusted-SE | z-value | p-value | IC2.5% | IC97.5% |
| --- | --- | --- | --- | --- | --- | --- | --- | --- |
| $N_{ppp}$ | Experimental group | 0.01 | 0.05 | 0.05 | 0.22 | 0.830 | -0.15 | 0.27 |
|  | Sex | -0.12 | 0.12 | 0.12 | 0.97 | 0.332 | -0.39 | 0.03 |
| $\Sigma[\text{pesticide}]_{scaled}$ | Sex | -0.35 | 1.28 | 1.30 | 0.27 | 0.785 | -6.11 | 3.06 |
|  | BC | 0.25 | 1.20 | 1.22 | 0.23 | 0.819 | -3.33 | 5.90 |
| $TI_{tot} (\times 10^{-3})$ | Experimental group | -0.20 | 0.20 | 0.20 | 0.97 | 0.331 | -0.60 | 0.03 |
|  | Sex | 0.10 | 0.10 | 0.10 | 0.62 | 0.536 | -0.10 | 0.50 |

**Figure 3:**
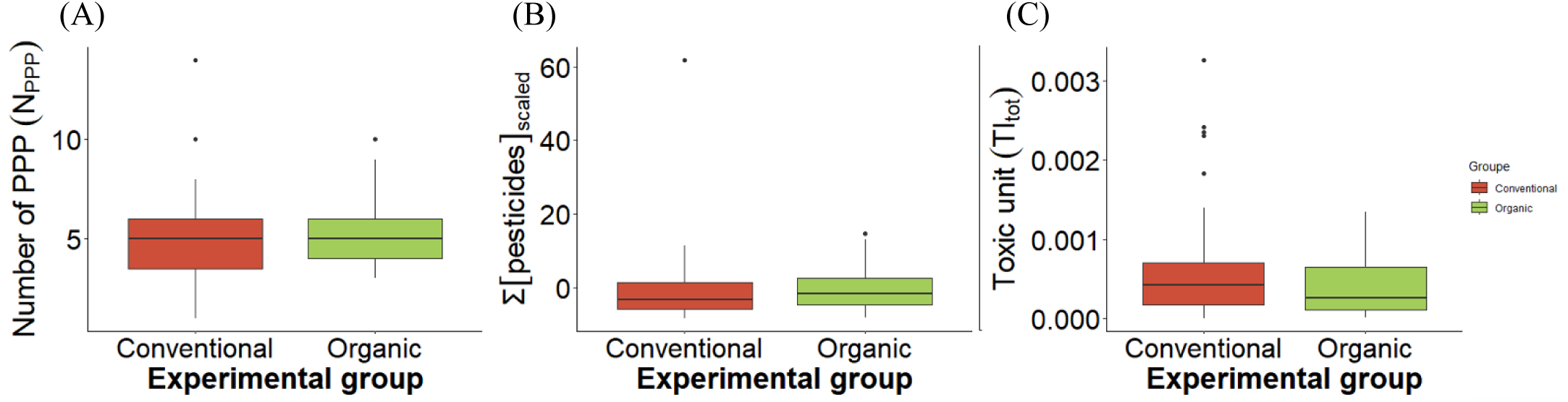
Effect of experimental food treatment (conventional, in red *vs* organic, in green) on (A) the number of PPPs, (B) the cumulative sum of scaled pesticides levels and (C) the global toxic unit. Box-and whisker plots represent the data: the top and bottom of the boxes represent the first and last quartiles, respectively; the line across the box represents the median. The whiskers represent the 5th and 95th percentiles, and the circles represent outliers. Significant difference is indicated by an asterisk (*p-value* < 0.050).

### (b) Food treatment dissimilarities in frequency and abundance

Multivariate dispersion (mean distance to group centroids) was similar across the two feeding treatments for both similarity indices (Jaccard distance: F_1,68_ = 1.06, *p* = 0.306; Bray–Curtis distance: F_1,68_ = 1.28, *p* = 0.262), indicating comparable variability within the conventional and organic groups. However, the two feeding groups differed slightly in both the set of pesticides detected and their concentrations in partridge blood, as indicated by PERMANOVA on Jaccard and Bray–Curtis distance matrices (Jaccard distance: F_1,68_ = 2.00, *p* = 0.049; Bray–Curtis distance: F_1,68_ = 2.36, *p* = 0.040). As examples, Nitenpyram was detected in a higher proportion in conventionally-than organically-fed individuals (31 *vs* 17, respectively, Chi-square test: χ² = 10.37, *p* = 0.001, Table 2), while the reverse was observed for Bifenox, Dimethenamid-P, Diflufenican and Epoxiconazole (20 *vs* 28, χ² = 4.05, *p* = 0.044; 4 *vs* 12, F = 4.14, *p* = 0.024; 3 *vs* 10, F = 4.35, *p* = 0.034 and 1 *vs* 8, F = 10.16, *p* = 0.013, respectively, Table 2). Moreover, Bifenox was quantified at higher concentrations in conventionally-fed birds (i.e., conventional *vs* organic: 1644.36 ± 374.60 mg.kg^-1^ *vs* 818.21 ± 152.24 mg.kg^-1^, W = 388.00, *p* = 0.024), while Dimethenamid-P was quantified at higher concentrations in organically-fed partridges (i.e., conventional vs organic: 0.03 ± 0.01 mg.kg^-1^ *vs* 0.19 ± 0.04 mg.kg^-1^, W = 7.00, *p* = 0.042).

**Figure 4:**
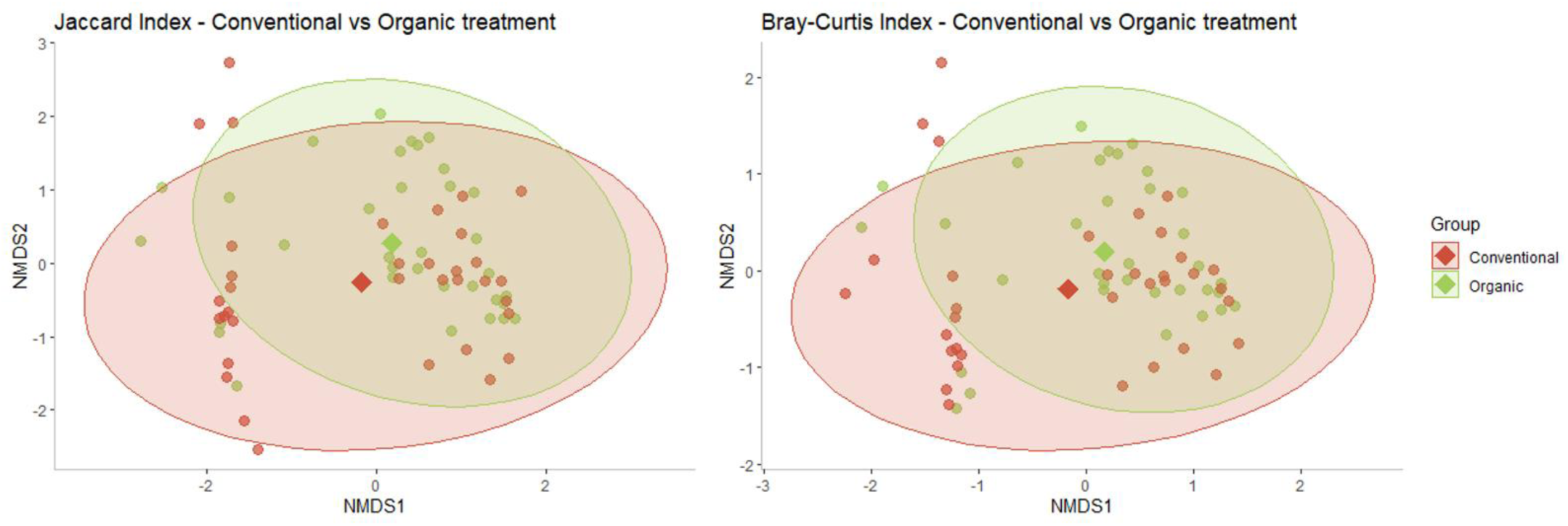
Nonmetric multidimensional scaling (NMDS) ordination in two-dimensional space of (A) set of pesticides an individual is contaminated by, calculated through the Jaccard index, and (B) PPP levels, estimated using the Bray-Curtis index, for the 80 semi-captive partridges studied here. As each dot represents one individual, NMDS plots help here visualize individual-level divergence and relatedness between food treatments. Red and green ellipses indicate the distribution of conventionally- and organically-fed birds respectively. The red and green diamonds correspond to centroid for conventionally- and organically-fed individuals respectively.

## 4. DISCUSSION

This study aimed to assess the importance of food ingestion in routes of contamination by evaluating differences in PPP patterns between semi-captive partridges fed conventional grains and those fed organic grains over a five-month period. Contrary to our first prediction, the most striking result of our study was that organically-fed partridges, despite being fed grains with no PPP quantified, showed similar levels and diversity of PPPs detected in their blood to those observed in conventionally-fed birds. Such a result tends to indicate that food may not necessarily be the only source of contamination for non-target organisms. Moreover, organic feeding alone may not be sufficient to prevent PPP contamination. However, marginal distinctions in PPP contamination profiles can be drawn between our two food treatments, with certain molecules appearing to be more abundant and/or at higher concentrations in either the conventional or organic food treatment. Overall, these findings highlight the concerning multiple and widespread environmental sources of PPP contamination, with absorptive food impregnation playing possibly only a negligible role.

Firstly, the weak connections among compounds applied on crops, residues present in commercial juvenile poultry food, and conventional grains, and those found in both conventionally- and organically-fed partridge blood cast doubt on the ingestion exposure hypothesis as the primary route of contamination. Indeed, less than half of the number of compounds quantified in the juvenile pellet and the experimental feed mixture (i.e., Piperonyl butoxyde and Tebuconazole) were detected in the blood of a few partridges from both experimental groups (conventional and organic, see Table 3). Such a discrepancy between food PPP content and partridge blood PPP profiles might initially be explained by the striking difference in PPP load measurement sensibility for food and blood. Indeed, the PPP quantification technique applied to foods was six times less sensitive than that used for blood. It means that we might have missed part of the PPPs that the partridges were exposed to through the food provided. Additionally, although not detected in the provided foods (due to limited sensitivity), the compounds contaminating them might be detected and quantified in partridge blood due to their bioaccumulation during the five-month experimental exposure. Nevertheless, such an ultimate hypothesis must be confirmed, as we monitored partridge contamination through blood analyses, which usually reflect short-term organismal PPP activity. To note, we might have missed part of the partridge PPP contamination profile because the analytical techniques used on blood matrix are biased toward hydrophilic compounds (Rodrigues et al., 2023). Indeed, certain substances might have lipophilia affinity which induce their rapid storage in fat or feather, some may also be accumulated in specific organs (such as liver), and finally others may be efficiently excreted and were then not found in the studied blood matrix (Espin et al. 2016; Pacyna-Kuchta, 2023). To get a complete overview of partridges’ contamination, the next step would be to measure PPPs in two complementary biological minimally-invasive matrixes such as blood and feather. Overall, the absence of congruence between PPP in conventional grains and partridge PPP contamination profiles found in this study strongly suggests that bird exposure to pesticides is not mainly due to their food and arises from the environment (i.e., air, soil and/or water). In other words, in the global current context of pervasive PPP exposure, atmospheric deposition, contaminated drinking water and persistence-driven remobilization from soils likely act in combination, resulting in continuous multi-route exposure (i.e., respiratory pathways, during preening, and/or dermal exposure) that overrides the food contamination implemented in this study (Mineau 2011; Quaglia et al. 2019; Sánchez-Bayo 2011). Future research should consider using techniques such as air analysis (e.g., using passive air samplers or active air sampling devices), analysis of water consumed by studied non-target organisms), as well as soil sampling (e.g., core samples from the organisms’ housing environment) to better understand the pathways through which non-target organisms have been exposed.

Off-field transport from surrounding conventional agricultural areas (which represent 99.02% of agricultural areas within a 1-km radius of the two pens) is a plausible source. Spray drift and volatilization followed by atmospheric transport and redeposition through rainfall are well-documented processes that can contaminate non-target environments several hundred kilometers away from treated fields (Shen et al. 2005; Soccoro et al. 2016). In that context, PPPs applied on both neighboring and at-distance crops may have reached the experimental pens through air masses and subsequent wet deposition, independently of the birds’ diet. Consistently, data from the French national pesticide sales database (BNV-d) indicate that, locally, Pendimethanil, Dimethenamid-P and Diflufenican were among the most sold active substances (after glyphosate) within the postal codes surrounding the experimental site in 2020. These molecules were detected in partridge blood at frequencies of 24.64%, 23.18%, and 18.84%, respectively, supporting the hypothesis that local agricultural use is a significant driver of contamination. Moreover, Diphenylamine, one of the most frequently detected PPPs in our partridges, is a widely used growth regulator with fungicidal properties, applied globally to control post-harvest diseases in fruits, and is often detected as residues in soils worldwide (EFSA 2012).

Regarding a potential soil-borne exposure pathway, it is important to note that the soil within the aviaries has not been subjected to any pesticide treatment since the establishment of the commercial game farm, i.e., for more than two decades. In such a context, the presence of recently applied compounds appears unlikely. In that context, only persistent active substances would be expected to remain detectable through remobilization, runoff, or resuspension of contaminated particles, leading to chronic low-level exposure *via* inhalation of dust. This may notably apply to compounds such as Diphenylamine and Epoxiconazole which were recognized for their marked persistence in French agricultural soils (Froger et al. 2023).

Water also represents a credible exposure route, both for approved and banned active substances detected in partridge blood. First, the drinking water supplied to both conventionally- and organically-fed partridges was collected from the public water network intended for human consumption. Although treated, such water may still contain trace levels of PPPs, especially in agricultural regions where water contamination is strongly correlated with agricultural practices (and compounds purchases by postcode notably, Rigal & Perrot, 2025; Staub et al. 2025). However, since this water is intended for human consumption, such a source of contamination seems unlikely, given the very strict regulatory thresholds for acceptable contamination in place. Another hypothesis of water-originated contamination comes from water run-off, especially in agricultural landscape. Accordingly, surface water monitoring data available from the Naiades database reported the presence of several substances that were detected in partridge blood in Deux-Sèvres’s sampling stations (N = 66). As an example, Bifenox, the third most frequently detected PPP in partridges’ blood was sampled in February and March 2021 in more than 15 different stations around the game farm. Additionally, Diphenylamine, Tolylfluanid and Carbendazim, three of the eight compounds with the highest detection frequency in partridge blood, are extensively persistent in water. Despite being banned by EC Regulation 1107/2009 for up to 10 years (i.e., 18, 14 and 12 years respectively; Singer et al. 2010; Reemtsma et al. 2013), they were sampled in 10, 3 and 20 stations in the Deux-Sèvres department in March 2021 respectively. Altogether, this spatial concordance supports the likelihood of water run-off exposure.

Interestingly, although the overall PPP contamination pattern did not differ in the number of compounds or concentrations between the two food treatments, we observed PPP contamination frequency and abundance dissimilarities between conventionally-fed and organically-fed birds. Furthermore, we highlighted a greater (but non-significant) variability among conventionally-than organically-fed birds. These results are particularly surprising because the two aviaries were next to each other and contained the exact same equipment. Several non-mutually exclusive hypotheses can explain these results. First, while adjacent, the two aviaries might present some micro-environmental differences: the conventional aviary was closer to the proximate conventional agricultural area (and no hedge was buffering possible PPP transported through air), while the organic group aviary was more submitted to West wind, i.e., air and dust dispersion, than the conventional group. However, it is unlikely that such minor abiotic differences could explain the contamination differences observed, considering the very close proximity of the two aviaries. Secondly, the contamination differences might not originate from abiotic conditions but be explained by some biotic features, such as differential individual detoxication capacities. Indeed, avian species exhibit varying metabolic rates, which can impact their ability to metabolize and eliminate PPPs from their bodies (Fuentes et al., 2024; Katagi & Fujisawa, 2021; Mineau, 2011). As a result, any differences in PPP contamination levels between food treatments may have been obscured due to variations in metabolism and elimination rates among partridges. The differences we observed between conventionally- and organically-fed individuals could notably be mediated by divergent microbiome composition and function but this hypothesis should be properly investigated. On that line, blood PPP profile may reflect buffered PPP contamination rather than the entire PPP environmental exposure.

Overall, the start of our experiment coincided with both food change (i.e., transition from juvenile pellets to seed mixtures) and seasonal farming practices, i.e., summer crop applications *vs* autumn/winter treatments for overwintering and soil preparation. As a result, birds faced both potential microbiota changes and novel environmental PPPs, supporting that variations in contamination profiles are driven by both food and environmental factors (Bariod et al., 2025b; Crisol-Martinez et al., 2016; Ruuskanen et al., 2020; Sun et al., 2022).

Finally, the influence of individual traits on PPP contamination in blood could help explain the differential contamination levels among individuals. Grey partridges exhibit a wide range of known physiological mechanisms that can response to PPP contamination, such as fat stores, hormone levels or immunological pathways (e.g., Gaffard et al 2022a and b, Moreau et al. 2021 and 2022a, see Moreau et al. 2022b for a review). For instance, partridges with higher detoxification capacities could show higher concentrations of molecules involved in immunological responses (e.g., carotenoids pigments; Lopez-Antia et al. 2013, 2015a, b or haptoglobin protein; Quaye 2008) to mitigate PPP-induced oxidative stress. Examining hormonal differences, metabolic rates and behaviors could provide insights into these contamination patterns.

## 5. CONCLUSION

Here, the large-scale quantification and variability of PPPs in semi-captive partridge blood revealed their widespread presence in blood, whatever the food treatment (i.e., conventional or organic). This pattern suggests that food ingestion may play only a marginal role as a source of contamination for non-target vertebrates inhabiting agroecosystems, with exposure instead driven by continuous, multi-route pathways. Indeed, any differences are largely overwhelmed by a pesticide-saturated environment that affects individuals in a largely indiscriminate manner.

Moreover, both abiotic and biotic factors such as PPP persistence in the environment, metabolism or microbiota are likely contributors.

Importantly, our study was only focused on food contamination, overlooking the broader ecosystemic benefits of organic farming (with only 0.98% of agricultural areas within a 1-km radius around pens under organic practices). This reinforces that assessing organic food alone, i.e., without accounting for landscape dynamics and ecosystem services notably, fails to capture the full benefits of organic farming systems. Yet, monitoring PPP contamination across trophic levels, and especially in humans, consuming non-target organisms living in agricultural landscape, should be integrated into a One Health framework in a close future (Xie et al., 2017).

## Supporting information

Script analyses

## ACKNOWLEDGEMENTS

We are indebted to thank Julien Blandin for letting us work on his farm and for looking after the partridges all along the experiment. We also would like to thank the Fédération des Chasseurs (79), its president Guy Guédon, Frédéric Audurier, and their team for their support. We are grateful to Landry Boussac and Simon Billon for their help in birds monitoring.

## Appendix 1: Composition, analytical constituents, and additives found in ground-feed mixture “STARGIB entretien” provided *ad libitum* to grey partridges from birth to the start of the experiment.

Composition: wheat, wheat bran, wheat middlings, shelled sunflower meal, soybean meal, corn, barley, rapeseed meal, sunflower meal, fruit pomace, calcium carbonate, dicalcium phosphate, cane molasses, additive premixes, salt, sodium bicarbonate

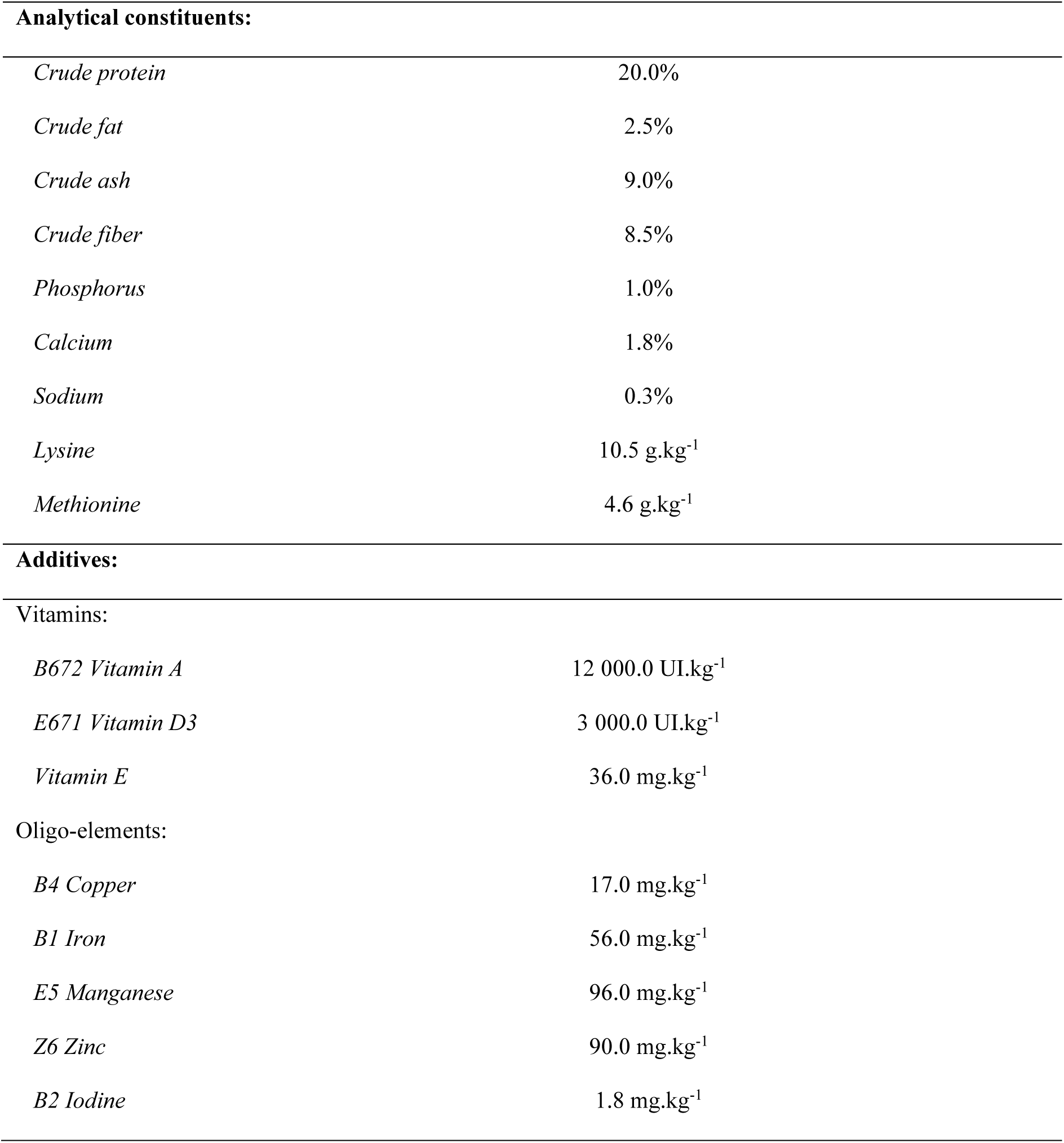
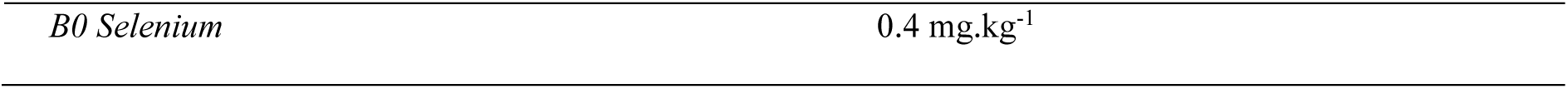

## Appendix 2 Lists of phytosanitary treatments applied by farmers to crops that produced corn, pea, wheat and faba bean provided to conventionally-fed partridges during the experimentation (from the 30^th^ of October 2020 to the 24^th^ of March 2021).

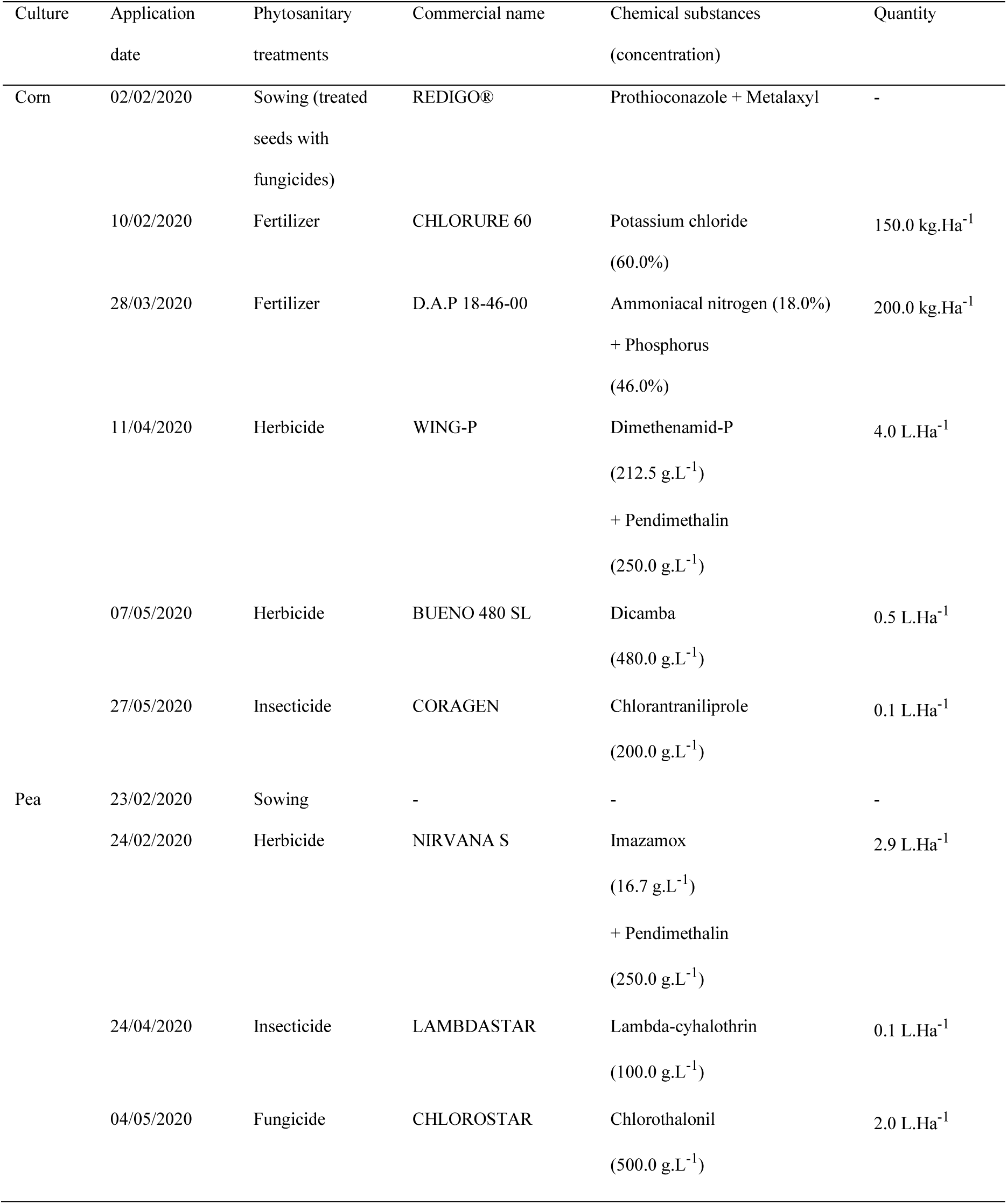
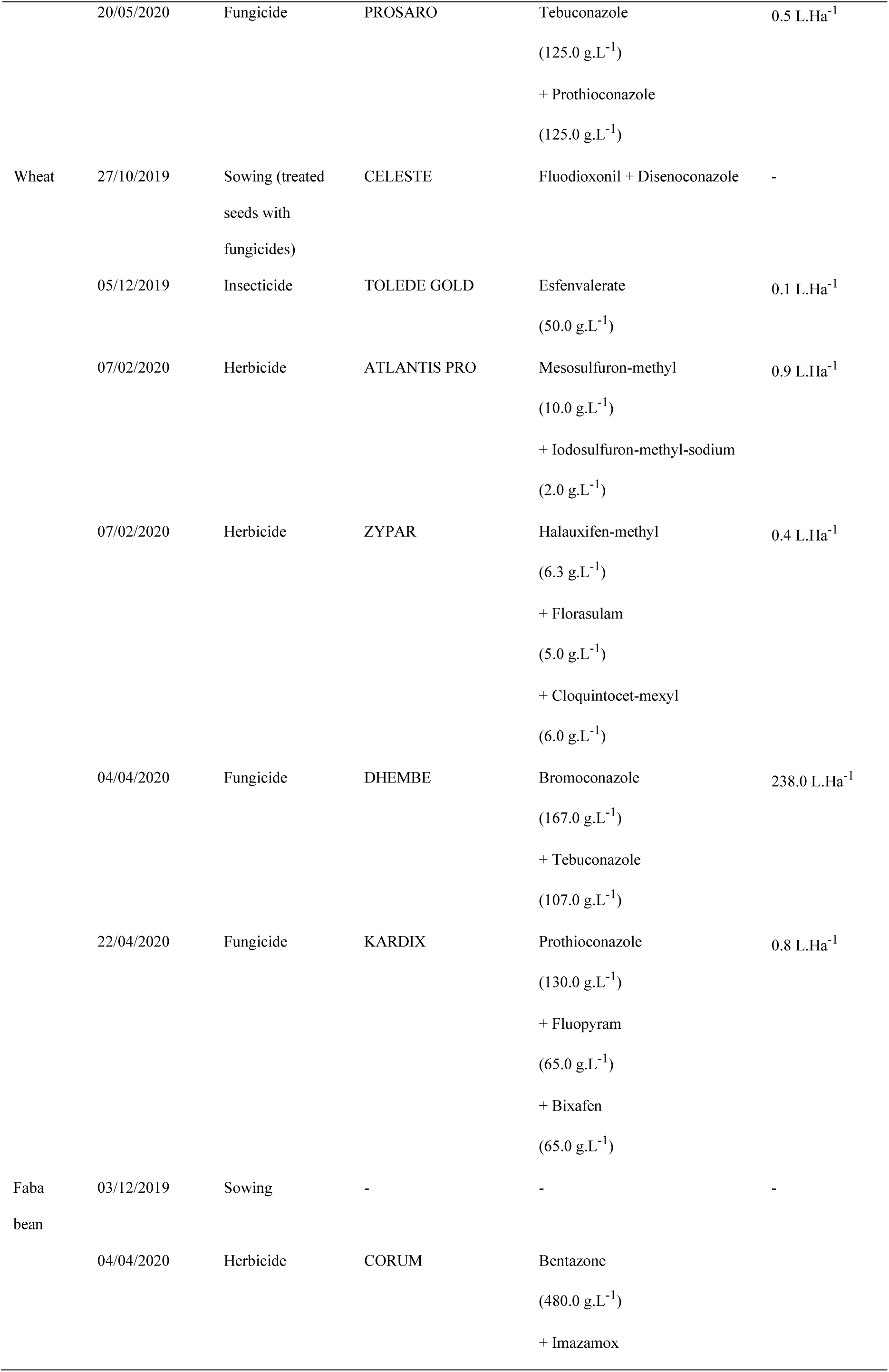
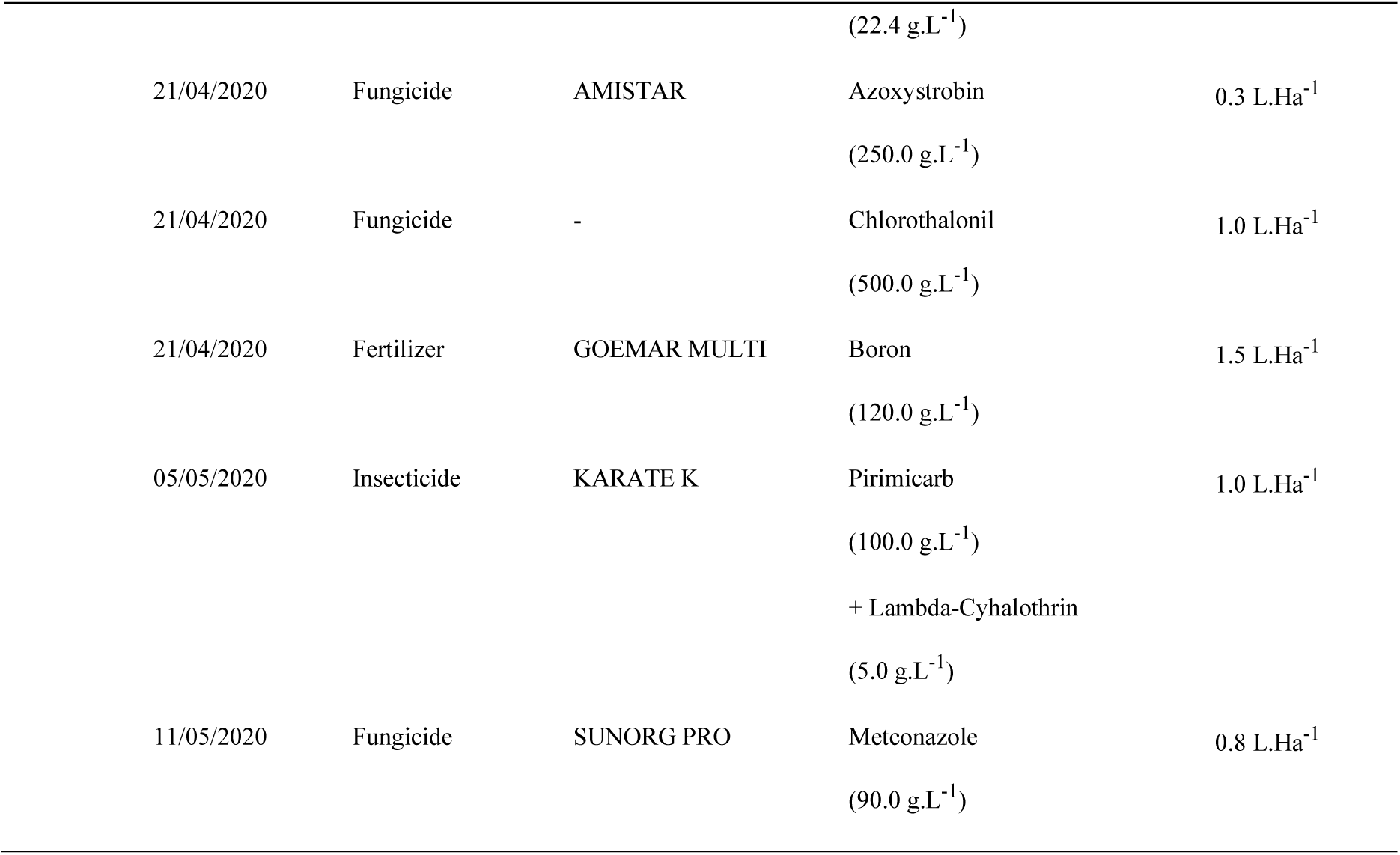

## Appendix 3: Characteristics of the 33 PPPs detected in 69 partridge blood samples ordered by decreasing number of occurrences (>LOD). Lab species for Bird LD50 values are *Colinus virginianus*, *Coturnix. c. japonica*, *Anas platyrhynchos*, *Phasianus sp.* and *Serinus canaria*. European Commission Regulation 1107/2009 status. Farm means farming use; Dom means domestic use. *** not a PPP, may be authorized at national level under different legislation. Modes of Action came from Insecticide, Fungicide and Herbicide Resistance Action Committee (IRAC, FRAC, HRAC)

| Name | Type | Chemical family | Use | EC regulation | Expiration<br>of<br>approval | Bird<br>LD50<br>(mg.kg<sup>-1</sup>) | LOD<br>× 10<sup>-3</sup><br>(mg.kg<sup>-1</sup>) | LOQ<br>× 10<sup>-3</sup><br>(mg.kg<sup>-1</sup>) | Detected<br>samples | Mode of Action |
| --- | --- | --- | --- | --- | --- | --- | --- | --- | --- | --- |
| Diphenylamine | Fungicide | Amine | Farm | disapproved | 2012 | 2250.0 | 0.001 | 0.004 | 55 | Polyene and isoprene biosynthesis inhibitor |
| Nitenpyram | Insecticide | Neonicotinoid | Dom | *** | - | 1124.0 | 0.009 | 0.030 | 48 | Nicotinic acetylcholine receptor competitive modulator |
| Bifenox | Herbicide | Diphenyl-ether | Farm | approved | - | 2000.0 | 0.001 | 0.004 | 48 | Inhibition of protoporphyrinogen oxidase |
| Tolylfluanid | Fungicide | Sulfonamide | Farm | disapproved | 2007 | 2000.0 | 0.008 | 0.025 | 38 | Multi-site activity with protective action |
| Pendimethalin | Herbicide | Dinitroanilin | Farm | approved | - | 1421.0 | 0.025 | 0.083 | 17 | Inhibition of microtubule assembly |
| Dimethenamid-P | Herbicide | Chloroacetamide | Farm | approved | - | 1068.0 | 0.028 | 0.094 | 16 | Inhibition of cell division |
| Diflufenican | Herbicide | Carboxamide | Farm | approved | - | 2150.0 | 0.003 | 0.010 | 13 | Inhibition of phytoene desaturase |
| Carbendazim | Fungicide | Benzimidazole | Dom | disapproved | 2014 | 2250.0 | 0.004 | 0.014 | 11 | Cytoskeleton and motor protein |
| Cyprodinil | Fungicide | Anilino-pyrimidine | Farm | approved | - | 500.0 | 0.001 | 0.004 | 11 | Amino acids and protein synthesis |
| Epoxiconazole | Fungicide | Triazole | Farm | disapproved | 2020 | 2000.0 | 0.003 | 0.009 | 9 | Sterol biosynthesis in membranes inhibitor |
| Propiconazole | Fungicide | Triazole | Farm | disapproved | 2018 | 2510.0 | 0.001 | 0.003 | 8 | Sterol biosynthesis in membranes inhibitor |
| Quinoxifen | Fungicide | Quinolein | Farm | disapproved | 2019 | 2250.0 | 0.005 | 0.016 | 8 | Signal transduction |
| Thiametoxam | Insecticide | Neonicotinoid | Farm | disapproved | 2018 | 576.0 | 0.012 | 0.041 | 8 | Nicotinic acetylcholine receptor competitive modulator |
| Alachlor | Herbicide | Chloroacetanilide | Farm | disapproved | 2006 | 1536.0 | 0.028 | 0.093 | 7 | Elongase and geranylgeranyl pyrophosphate cyclisation enzymes inhibition |
| Flusilazole | Fungicide | Triazole | Farm | disapproved | 2008 | 1590.0 | 0.014 | 0.048 | 7 | Sterol biosynthesis in membranes inhibitor |
| Penconazole | Fungicide | Triazole | Farm | approved | - | 1590.0 | 0.001 | 0.003 | 5 | Sterol biosynthesis in membranes inhibitor |
| Myclobutanil | Fungicide | Triazole | Farm, Dom | disapproved | 2021 | 510.0 | 0.021 | 0.071 | 5 | Sterol biosynthesis in membranes inhibitor |
| Propyzamid | Herbicide | Benzonitrile | Farm | approved | - | 6578.0 | 0.002 | 0.007 | 5 | Inhibition of microtubule assembly |
| Metolachlor-S | Fungicide | Chloroacetanilide | Farm | approved | - | 2000.0 | 0.010 | 0.032 | 5 | Elongase and geranylgeranyl pyrophosphate cyclisation enzymes inhibition |
| Triflusaluron-methyl | Herbicide | Sulfonylurea | Farm | approved | - | 2250.0 | 0.001 | 0.004 | 4 | Inhibition of acetolactate synthase |
| Clomazone | Herbicide | Isoxazolidinone | Farm | approved | - | 2224.0 | 0.012 | 0.040 | 3 | Inhibition of deoxy-d-xyulose phosphate synthase |
| Piperonyl butoxide | Insecticide | Benzodioxol | Farm, Dom | *** | - | 2250.0 | 0.000 | 0.001 | 2 | Cytochrome P450s inhibitor |
| Tebuconazole | Fungicide | Triazole | Farm | approved | - | 1988.0 | 0.001 | 0.005 | 2 | Sterol biosynthesis in membranes inhibitor |
| Carbetamide | Herbicide | Carbamates | Farm, Dom | disapproved | 2021 | 2000.0 | 0.005 | 0.018 | 2 | Inhibition of microtubule organisation |
| Pyrimethanil | Fungicide | Pyridine | Farm | approved | - | 2000.0 | 0.150 | 0.500 | 2 | Amino acids and protein synthesis |
| Trifluralin | Herbicide | Dinitroaniline | Farm | disapproved | 2010 | 2250.0 | 0.008 | 0.028 | 2 | Inhibition of microtubule assembly and cell division inhibitor |
| Dichlorprop-methyl | Herbicide | Phenoxy-carboxylates | Farm | approved | - | 4640.0 | 0.018 | 0.061 | 1 | Active auxin synthetic promoter |
| Spinosad-A | Insecticide | Macrolid | Farm, Dom | approved | - | 1333.0 | 0.000 | 0.001 | 1 | Nicotinic acetylcholine receptor competitive modulator |
| Dimethachlor | Herbicide | Chloroacetanilide | Farm | approved | - | 524.0 | 0.028 | 0.094 | 1 | Elongase and geranylgeranyl pyrophosphate cyclisation enzymes inhibition |
| Fenpropidin | Fungicide | Piperidin | Farm | approved | - | 1899.0 | 0.008 | 0.025 | 1 | Sterol biosynthesis in membranes inhibitor |
| Fenpropimorph | Fungicide | Morpholin | Farm | disapproved | 2019 | 2000.0 | 0.003 | 0.009 | 1 | Sterol biosynthesis in membranes inhibitor |
| Isoxaflutole | Herbicide | Isoxazole | Farm | approved | - | 2150.0 | 0.038 | 0.128 | 1 | Inhibition of hydroxyphenyl pyruvate dioxygenase |
| Thiacloprid | Insecticide | Neonicotinoid | Farm | disapproved | 2018 | 35.0 | 0.001 | 0.005 | 1 | Nicotinic acetylcholine receptor competitive modulator |

