## Supplementary material for "High residual pesticide contamination despite contrasted feeding treatments in semi-captive bred grey partridges": Script analyses

rm(list=ls())

\## Library

library(pavo)

library(ade4)

library(magrittr)

library(factoextra)

library(maptools)

library(gplots)

library(nlme)

library(lme4)

library(lmerTest)

library(ggplot2)

library(dplyr)

library(lubridate)

library(gghalves)

library(car)

library(lsmeans)

library(MuMIn)

library(Hmisc)

library(multcomp)

library(effects)

library(phia)

library(FactoMineR)

library(mgcv)

library(languageR)

library(knitr)

library(rstatix)

library(tidyverse)

library(vegan)

library(RVAideMemoire)

library(reshape2)

library(devtools)

library(pairwiseAdonis)

library(rptR)

library(effectsize)

library(arm)

library(smatr)

library(cowplot)

library(corrplot)

#Opening dataset

setwd("C:/")

data <- read.csv("Data_Partridge_analyses_total.csv", sep=";", h=T)

#Creating relevant working subsets

conventional = subset(data, data$Experimental_group=="Conventional")

organic = subset(data, data$Experimental_group=="Organic")

#Standard error calculation

std <- function(x) sd(x)/sqrt(length(x))

###Pearson correlation test between the three PPP loads of interest

cor.test(data$Nppp, data$sum_pesticides_scale, method="pearson")

cor.test(data$Nppp, data$TI_tot, method="pearson")

cor.test(data$TI_tot, data$sum_pesticides_scale, method="pearson")

\##### Figure 2, Heatmap

str(data)

heatmap <- melt(setDT(data), id.vars = c("Partridge_ID"), variable.name = "mol")

df_heatmap <- ddply(data, .(mol), transform,

rescale = scale(value))

df_heatmap <- rename.variable(df_heatmap, "Partridge_ID", "Individuals")

df_heatmap <- rename.variable(df_heatmap, "mol", "Compounds")

df_heatmap <- rename.variable(df_heatmap, "value", "Concentration")

df_heatmap$Concentration <- ifelse(df_heatmap$Concentration > 0,

df_heatmap$Concentration, NA)

ggplot(df_heatmap, aes(Individuals, Compounds, fill= Concentration)) +

geom_tile() +

scale_fill_gradient(low = "grey85", high = "black", na.value = "white")+

scale_x_discrete(position="top", expand = c(0,0)) +

scale_y_discrete(limits = rev(levels(df_heatmap$Compounds)))+

theme(legend.title = element_text(color="black", size="15")) +

theme(legend.text = element_text(color="black", size="15")) +

theme(axis.text.x= NULL)+

theme(axis.text.y=element_text(color="black", size="15"))+

labs(y=NULL,x=NULL)

####Fisher and wilcoxon test (Table 2)

fisher <- read.csv("Fishertest_Patridge.csv", sep=";", h=T)

quantity <- data

\## Alachlor

Alachlor_descriptive <- quantity %>%

filter(Alachlor > 0)

Alachlor_descriptive %>%

group_by(Experimental_group) %>%

summarise(

no_rows = length(Alachlor),

average = mean(Alachlor, na.rm = TRUE),

stderror = std(Alachlor),

median	= median(Alachlor, na.rm = TRUE),

Q1	= quantile(Alachlor, 0.25, na.rm = TRUE),

Q3	= quantile(Alachlor, 0.75, na.rm = TRUE),

min	= min(Alachlor, na.rm = TRUE),

max	= max(Alachlor, na.rm = TRUE),

.groups = "drop"

) %>%

mutate(across(where(is.numeric),

~ formatC(.x, digits = 6, format = "f")))

\## Bifenox

Bifenox_presence <- fisher[,c(3,5)]

Bifenox_quantity <- quantity[,c(3,5)]

chisq.test(table(Bifenox_presence))

Bifenox_detected <- Bifenox_presence %>%

filter(Bifenox > 0)

Bifenox_detected %>%

group_by(Groupe) %>%

summarise(no_rows = length(Bifenox))

Bifenox_quantified <- Bifenox_quantity %>%

filter(Bifenox > 0)

wilcox.test(Bifenox ~ Experimental_group, data=Bifenox_quantified)

Bifenox_quantified %>%

group_by(Experimental_group) %>%

summarise(

no_rows = length(Bifenox),

average = mean(Bifenox, na.rm = TRUE),

stderror = std(Bifenox),

median	= median(Bifenox, na.rm = TRUE),

Q1	= quantile(Bifenox, 0.25, na.rm = TRUE),

Q3	= quantile(Bifenox, 0.75, na.rm = TRUE),

min	= min(Bifenox, na.rm = TRUE),

max	= max(Bifenox, na.rm = TRUE),

.groups = "drop"

) %>%

mutate(across(where(is.numeric),

~ formatC(.x, digits = 6, format = "f")))

\## Carbendazim

Carbendazim_presence <- fisher[,c(3,6)]

Carbendazim_quantity <- quantity[,c(3,6)]

chisq.test(table(Carbendazim_presence))

Carbendazim_detected <- Carbendazim_presence %>%

filter(Carbendazim > 0)

Carbendazim_detected %>%

group_by(Experimental_group) %>%

summarise(no_rows = length(Carbendazim))

Carbendazim_quantified <- Carbendazim_quantity %>%

filter(Carbendazim > 0)

wilcox.test(Carbendazim ~ Experimental_group, data=Carbendazim_quantified)

Carbendazim_quantified %>%

group_by(Experimental_group) %>%

summarise(

no_rows = length(Carbendazim),

average = mean(Carbendazim, na.rm = TRUE),

stderror = std(Carbendazim),

median	= median(Carbendazim, na.rm = TRUE),

Q1	= quantile(Carbendazim, 0.25, na.rm = TRUE),

Q3	= quantile(Carbendazim, 0.75, na.rm = TRUE),

min	= min(Carbendazim, na.rm = TRUE),

max	= max(Carbendazim, na.rm = TRUE),

.groups = "drop"

) %>%

mutate(across(where(is.numeric),

~ formatC(.x, digits = 6, format = "f")))

\## Carbetamide

Carbetamide_descriptive <- quantity %>%

filter(Carbetamide > 0)

Carbetamide_descriptive %>%

group_by(Experimental_group) %>%

summarise(

no_rows = length(Carbetamide),

average = mean(Carbetamide, na.rm = TRUE),

stderror = std(Carbetamide),

median	= median(Carbetamide, na.rm = TRUE),

Q1	= quantile(Carbetamide, 0.25, na.rm = TRUE),

Q3	= quantile(Carbetamide, 0.75, na.rm = TRUE),

min	= min(Carbetamide, na.rm = TRUE),

max	= max(Carbetamide, na.rm = TRUE),

.groups = "drop"

) %>%

mutate(across(where(is.numeric),

~ formatC(.x, digits = 6, format = "f")))

\## Clomazone

Clomazone_descriptive <- quantity %>%

filter(Clomazone > 0)

Clomazone_descriptive %>%

group_by(Experimental_group) %>%

summarise(

no_rows = length(Clomazone),

average = mean(Clomazone, na.rm = TRUE),

stderror = std(Clomazone),

median	= median(Clomazone, na.rm = TRUE),

Q1	= quantile(Clomazone, 0.25, na.rm = TRUE),

Q3	= quantile(Clomazone, 0.75, na.rm = TRUE),

min	= min(Clomazone, na.rm = TRUE),

max	= max(Clomazone, na.rm = TRUE),

.groups = "drop"

) %>%

mutate(across(where(is.numeric),

~ formatC(.x, digits = 6, format = "f")))

\## Cyprodinil

Cyprodinil_presence <- fisher[,c(3,10)]

Cyprodinil_quantity <- quantity[,c(3,10)]

chisq.test(table(Cyprodinil_presence))

Cyprodinil_detected <- Cyprodinil_presence %>%

filter(Cyprodinil > 0)

Cyprodinil_detected %>%

group_by(Experimental_group) %>%

summarise(no_rows = length(Cyprodinil))

Cyprodinil_quantified <- Cyprodinil_quantity %>%

filter(Cyprodinil > 0)

wilcox.test(Cyprodinil ~ Experimental_group, data=Cyprodinil_quantified)

Cyprodinil_quantified %>%

group_by(Experimental_group) %>%

summarise(

no_rows = length(Cyprodinil),

average = mean(Cyprodinil, na.rm = TRUE),

stderror = std(Cyprodinil),

median	= median(Cyprodinil, na.rm = TRUE),

Q1	= quantile(Cyprodinil, 0.25, na.rm = TRUE),

Q3	= quantile(Cyprodinil, 0.75, na.rm = TRUE),

min	= min(Cyprodinil, na.rm = TRUE),

max	= max(Cyprodinil, na.rm = TRUE),

.groups = "drop"

) %>%

mutate(across(where(is.numeric),

~ formatC(.x, digits = 6, format = "f")))

\## Dichlorprop

Dichlorprop_methyl_descriptive <- quantity %>%

filter(Dichlorprop_methyl > 0)

Dichlorprop_methyl_descriptive %>%

group_by(Experimental_group) %>%

summarise(

no_rows = length(Dichlorprop_methyl),

average = mean(Dichlorprop_methyl, na.rm = TRUE),

stderror = std(Dichlorprop_methyl),

median	= median(Dichlorprop_methyl, na.rm = TRUE),

Q1	= quantile(Dichlorprop_methyl, 0.25, na.rm = TRUE),

Q3	= quantile(Dichlorprop_methyl, 0.75, na.rm = TRUE),

min	= min(Dichlorprop_methyl, na.rm = TRUE),

max	= max(Dichlorprop_methyl, na.rm = TRUE),

.groups = "drop"

) %>%

mutate(across(where(is.numeric),

~ formatC(.x, digits = 6, format = "f")))

\## Diflufenican

Diflufenican_presence <- fisher[,c(3,13)]

Diflufenican_quantity <- quantity[,c(3,13)]

fisher.test(table(Diflufenican_presence))

Diflufenican_detected <- Diflufenican_presence %>%

filter(Diflufenican > 0)

Diflufenican_detected %>%

group_by(Experimental_group) %>%

summarise(no_rows = length(Diflufenican))

Diflufenican_quantified <- Diflufenican_quantity %>%

filter(Diflufenican > 0)

wilcox.test(Diflufenican ~ Experimental_group, data=Diflufenican_quantified)

Diflufenican_quantified %>%

group_by(Experimental_group) %>%

summarise(

no_rows = length(Diflufenican),

average = mean(Diflufenican, na.rm = TRUE),

stderror = std(Diflufenican),

median	= median(Diflufenican, na.rm = TRUE),

Q1	= quantile(Diflufenican, 0.25, na.rm = TRUE),

Q3	= quantile(Diflufenican, 0.75, na.rm = TRUE),

min	= min(Diflufenican, na.rm = TRUE),

max	= max(Diflufenican, na.rm = TRUE),

.groups = "drop"

) %>%

mutate(across(where(is.numeric),

~ formatC(.x, digits = 6, format = "f")))

\## Dimethachlor

Dimethachlor_descriptive <- quantity %>%

filter(Dimethachlor > 0)

Dimethachlor_descriptive %>%

group_by(Experimental_group) %>%

summarise(

no_rows = length(Dimethachlor),

average = mean(Dimethachlor, na.rm = TRUE),

stderror = std(Dimethachlor),

median	= median(Dimethachlor, na.rm = TRUE),

Q1	= quantile(Dimethachlor, 0.25, na.rm = TRUE),

Q3	= quantile(Dimethachlor, 0.75, na.rm = TRUE),

min	= min(Dimethachlor, na.rm = TRUE),

max	= max(Dimethachlor, na.rm = TRUE),

.groups = "drop"

) %>%

mutate(across(where(is.numeric),

~ formatC(.x, digits = 6, format = "f")))

\## Dimethenamid-P

dimethenamidP_presence <- fisher[,c(3,15)]

dimethenamidP_quantity <- quantity[,c(3,15)]

fisher.test(table(dimethenamidP_presence))

dimethenamidP_detected <- dimethenamidP_presence %>%

filter(DimethenamidP > 0)

dimethenamidP_detected %>%

group_by(Experimental_group) %>%

summarise(no_rows = length(DimethenamidP))

dimethenamidP_quantified <- dimethenamidP_quantity %>%

filter(DimethenamidP > 0)

wilcox.test(DimethenamidP ~ Experimental_group, data=dimethenamidP_quantified)

dimethenamidP_quantified %>%

group_by(Experimental_group) %>%

summarise(

no_rows = length(DimethenamidP),

average = mean(DimethenamidP, na.rm = TRUE),

stderror = std(DimethenamidP),

median	= median(DimethenamidP, na.rm = TRUE),

Q1	= quantile(DimethenamidP, 0.25, na.rm = TRUE),

Q3	= quantile(DimethenamidP, 0.75, na.rm = TRUE),

min	= min(DimethenamidP, na.rm = TRUE),

max	= max(DimethenamidP, na.rm = TRUE),

.groups = "drop"

) %>%

mutate(across(where(is.numeric),

~ formatC(.x, digits = 6, format = "f")))

\## Diphenylamine

Diphenylamine_presence <- fisher[,c(3,16)]

Diphenylamine_quantity <- quantity[,c(3,16)]

chisq.test(table(Diphenylamine_presence))

Diphenylamine_detected <- Diphenylamine_presence %>%

filter(Diphenylamine > 0)

Diphenylamine_detected %>%

group_by(Experimental_group) %>%

summarise(no_rows = length(Diphenylamine))

Diphenylamine_quantified <- Diphenylamine_quantity %>%

filter(Diphenylamine > 0)

wilcox.test(Diphenylamine ~ Experimental_group, data=Diphenylamine_quantified)

Diphenylamine_quantified %>%

group_by(Experimental_group) %>%

summarise(

no_rows = length(Diphenylamine),

average = mean(Diphenylamine, na.rm = TRUE),

stderror = std(Diphenylamine),

median	= median(Diphenylamine, na.rm = TRUE),

Q1	= quantile(Diphenylamine, 0.25, na.rm = TRUE),

Q3	= quantile(Diphenylamine, 0.75, na.rm = TRUE),

min	= min(Diphenylamine, na.rm = TRUE),

max	= max(Diphenylamine, na.rm = TRUE),

.groups = "drop"

) %>%

mutate(across(where(is.numeric),

~ formatC(.x, digits = 6, format = "f")))

\## Epoxiconazole

Epoxiconazole_presence <- fisher[,c(3,17)]

Epoxiconazole_quantity <- quantity[,c(3,17)]

fisher.test(table(Epoxiconazole_presence))

Epoxiconazole_detected <- Epoxiconazole_presence %>%

filter(Epoxiconazole > 0)

Epoxiconazole_detected %>%

group_by(Experimental_group) %>%

summarise(no_rows = length(Epoxiconazole))

Epoxiconazole_quantified <- Epoxiconazole_quantity %>%

filter(Epoxiconazole > 0)

wilcox.test(Epoxiconazole ~ Experimental_group, data=Epoxiconazole_quantified)

Epoxiconazole_quantified %>%

group_by(Experimental_group) %>%

summarise(

no_rows = length(Epoxiconazole),

average = mean(Epoxiconazole, na.rm = TRUE),

stderror = std(Epoxiconazole),

median	= median(Epoxiconazole, na.rm = TRUE),

Q1	= quantile(Epoxiconazole, 0.25, na.rm = TRUE),

Q3	= quantile(Epoxiconazole, 0.75, na.rm = TRUE),

min	= min(Epoxiconazole, na.rm = TRUE),

max	= max(Epoxiconazole, na.rm = TRUE),

.groups = "drop"

) %>%

mutate(across(where(is.numeric),

~ formatC(.x, digits = 6, format = "f")))

\## Fenpropidin

Fenpropidin_descriptive <- quantity %>%

filter(Fenpropidin > 0)

Fenpropidin_descriptive %>%

group_by(Experimental_group) %>%

summarise(

no_rows = length(Fenpropidin),

average = mean(Fenpropidin, na.rm = TRUE),

stderror = std(Fenpropidin),

median	= median(Fenpropidin, na.rm = TRUE),

Q1	= quantile(Fenpropidin, 0.25, na.rm = TRUE),

Q3	= quantile(Fenpropidin, 0.75, na.rm = TRUE),

min	= min(Fenpropidin, na.rm = TRUE),

max	= max(Fenpropidin, na.rm = TRUE),

.groups = "drop"

) %>%

mutate(across(where(is.numeric),

~ formatC(.x, digits = 6, format = "f")))

\## Fenpropimorph

Fenpropimorph_descriptive <- quantity %>%

filter(Fenpropimorph > 0)

Fenpropimorph_descriptive %>%

group_by(Experimental_group) %>%

summarise(

no_rows = length(Fenpropimorph),

average = mean(Fenpropimorph, na.rm = TRUE),

stderror = std(Fenpropimorph),

median	= median(Fenpropimorph, na.rm = TRUE),

Q1	= quantile(Fenpropimorph, 0.25, na.rm = TRUE),

Q3	= quantile(Fenpropimorph, 0.75, na.rm = TRUE),

min	= min(Fenpropimorph, na.rm = TRUE),

max	= max(Fenpropimorph, na.rm = TRUE),

.groups = "drop"

) %>%

mutate(across(where(is.numeric),

~ formatC(.x, digits = 6, format = "f")))

\## Flusilazole

Flusilazole_descriptive <- quantity %>%

filter(Flusilazole > 0)

Flusilazole_descriptive %>%

group_by(Experimental_group) %>%

summarise(

no_rows = length(Flusilazole),

average = mean(Flusilazole, na.rm = TRUE),

stderror = std(Flusilazole),

median	= median(Flusilazole, na.rm = TRUE),

Q1	= quantile(Flusilazole, 0.25, na.rm = TRUE),

Q3	= quantile(Flusilazole, 0.75, na.rm = TRUE),

min	= min(Flusilazole, na.rm = TRUE),

max	= max(Flusilazole, na.rm = TRUE),

.groups = "drop"

) %>%

mutate(across(where(is.numeric),

~ formatC(.x, digits = 6, format = "f")))

\## Isoxaflutole

Isoxaflutole_descriptive <- quantity %>%

filter(Isoxaflutole > 0)

Isoxaflutole_descriptive %>%

group_by(Experimental_group) %>%

summarise(

no_rows = length(Isoxaflutole),

average = mean(Isoxaflutole, na.rm = TRUE),

stderror = std(Isoxaflutole),

median	= median(Isoxaflutole, na.rm = TRUE),

Q1	= quantile(Isoxaflutole, 0.25, na.rm = TRUE),

Q3	= quantile(Isoxaflutole, 0.75, na.rm = TRUE),

min	= min(Isoxaflutole, na.rm = TRUE),

max	= max(Isoxaflutole, na.rm = TRUE),

.groups = "drop"

) %>%

mutate(across(where(is.numeric),

~ formatC(.x, digits = 6, format = "f")))

\## Metolachlor-S

MetolachlorS_descriptive <- quantity %>%

filter(MetolachlorS > 0)

MetolachlorS_descriptive %>%

group_by(Experimental_group) %>%

summarise(

no_rows = length(MetolachlorS),

average = mean(MetolachlorS, na.rm = TRUE),

stderror = std(MetolachlorS),

median	= median(MetolachlorS, na.rm = TRUE),

Q1	= quantile(MetolachlorS, 0.25, na.rm = TRUE),

Q3	= quantile(MetolachlorS, 0.75, na.rm = TRUE),

min	= min(MetolachlorS, na.rm = TRUE),

max	= max(MetolachlorS, na.rm = TRUE),

.groups = "drop"

) %>%

mutate(across(where(is.numeric),

~ formatC(.x, digits = 6, format = "f")))

\## Myclobutanil

Myclobutanil_descriptive <- quantity %>%

filter(Myclobutanil > 0)

Myclobutanil_descriptive %>%

group_by(Experimental_group) %>%

summarise(

no_rows = length(Myclobutanil),

average = mean(Myclobutanil, na.rm = TRUE),

stderror = std(Myclobutanil),

median	= median(Myclobutanil, na.rm = TRUE),

Q1	= quantile(Myclobutanil, 0.25, na.rm = TRUE),

Q3	= quantile(Myclobutanil, 0.75, na.rm = TRUE),

min	= min(Myclobutanil, na.rm = TRUE),

max	= max(Myclobutanil, na.rm = TRUE),

.groups = "drop"

) %>%

mutate(across(where(is.numeric),

~ formatC(.x, digits = 6, format = "f")))

\## Nitenpyram

Nitenpyram_presence <- fisher[,c(3,24)]

Nitenpyram_quantity <- quantity[,c(3,24)]

chisq.test(table(Nitenpyram_quantity))

Nitenpyram_detected <- Nitenpyram_presence %>%

filter(Nitenpyram > 0)

Nitenpyram_detected %>%

group_by(Experimental_group) %>%

summarise(no_rows = length(Nitenpyram))

Nitenpyram_quantified <- Nitenpyram_quantity %>%

filter(Nitenpyram > 0)

wilcox.test(Nitenpyram ~ Experimental_group, data=Nitenpyram_quantified)

Nitenpyram_quantified %>%

group_by(Experimental_group) %>%

summarise(

no_rows = length(Nitenpyram),

average = mean(Nitenpyram, na.rm = TRUE),

stderror = std(Nitenpyram),

median	= median(Nitenpyram, na.rm = TRUE),

Q1	= quantile(Nitenpyram, 0.25, na.rm = TRUE),

Q3	= quantile(Nitenpyram, 0.75, na.rm = TRUE),

min	= min(Nitenpyram, na.rm = TRUE),

max	= max(Nitenpyram, na.rm = TRUE),

.groups = "drop"

) %>%

mutate(across(where(is.numeric),

~ formatC(.x, digits = 6, format = "f")))

\## Penconazole

Penconazole_descriptive <- quantity %>%

filter(Penconazole > 0)

Penconazole_descriptive %>%

group_by(Experimental_group) %>%

summarise(

no_rows = length(Penconazole),

average = mean(Penconazole, na.rm = TRUE),

stderror = std(Penconazole),

median	= median(Penconazole, na.rm = TRUE),

Q1	= quantile(Penconazole, 0.25, na.rm = TRUE),

Q3	= quantile(Penconazole, 0.75, na.rm = TRUE),

min	= min(Penconazole, na.rm = TRUE),

max	= max(Penconazole, na.rm = TRUE),

.groups = "drop"

) %>%

mutate(across(where(is.numeric),

~ formatC(.x, digits = 6, format = "f")))

\## Pendimethalin

Pendimethalin_presence <- fisher[,c(3,26)]

Pendimethalin_quantity <- quantity[,c(3,26)]

chisq.test(table(Pendimethalin_presence))

Pendimethalin_detected <- Pendimethalin_presence %>%

filter(Pendimethalin > 0)

Pendimethalin_detected %>%

group_by(Experimental_group) %>%

summarise(no_rows = length(Pendimethalin))

Pendimethalin_quantified <- Pendimethalin_quantity %>%

filter(Pendimethalin > 0)

wilcox.test(Pendimethalin ~ Experimental_group, data=quantity1)

Pendimethalin_quantified %>%

group_by(Experimental_group) %>%

summarise(

no_rows = length(Pendimethalin),

average = mean(Pendimethalin, na.rm = TRUE),

stderror = std(Pendimethalin),

median	= median(Pendimethalin, na.rm = TRUE),

Q1	= quantile(Pendimethalin, 0.25, na.rm = TRUE),

Q3	= quantile(Pendimethalin, 0.75, na.rm = TRUE),

min	= min(Pendimethalin, na.rm = TRUE),

max	= max(Pendimethalin, na.rm = TRUE),

.groups = "drop"

) %>%

mutate(across(where(is.numeric),

~ formatC(.x, digits = 6, format = "f")))

\## Piperonyl Butoxide

Piperonylbutoxide_descriptive <- quantity %>%

filter(Piperonyl_butoxide > 0)

Piperonylbutoxide_descriptive %>%

group_by(Experimental_group) %>%

summarise(

no_rows = length(Piperonyl_butoxide),

average = mean(Piperonyl_butoxide, na.rm = TRUE),

stderror = std(Piperonyl_butoxide),

median	= median(Piperonyl_butoxide, na.rm = TRUE),

Q1	= quantile(Piperonyl_butoxide, 0.25, na.rm = TRUE),

Q3	= quantile(Piperonyl_butoxide, 0.75, na.rm = TRUE),

min	= min(Piperonyl_butoxide, na.rm = TRUE),

max	= max(Piperonyl_butoxide, na.rm = TRUE),

.groups = "drop"

) %>%

mutate(across(where(is.numeric),

~ formatC(.x, digits = 6, format = "f")))

\## Propiconazole

Propiconazole_descriptive <- quantity %>%

filter(Propiconazole > 0)

Propiconazole_descriptive %>%

group_by(Experimental_group) %>%

summarise(

no_rows = length(Propiconazole),

average = mean(Propiconazole, na.rm = TRUE),

stderror = std(Propiconazole),

median	= median(Propiconazole, na.rm = TRUE),

Q1	= quantile(Propiconazole, 0.25, na.rm = TRUE),

Q3	= quantile(Propiconazole, 0.75, na.rm = TRUE),

min	= min(Propiconazole, na.rm = TRUE),

max	= max(Propiconazole, na.rm = TRUE),

.groups = "drop"

) %>%

mutate(across(where(is.numeric),

~ formatC(.x, digits = 6, format = "f")))

\## Propyzamide

Propyzamide_descriptive <- quantity %>%

filter(Propyzamide > 0)

Propyzamide_descriptive %>%

group_by(Experimental_group) %>%

summarise(

no_rows = length(Propyzamide),

average = mean(Propyzamide, na.rm = TRUE),

stderror = std(Propyzamide),

median	= median(Propyzamide, na.rm = TRUE),

Q1	= quantile(Propyzamide, 0.25, na.rm = TRUE),

Q3	= quantile(Propyzamide, 0.75, na.rm = TRUE),

min	= min(Propyzamide, na.rm = TRUE),

max	= max(Propyzamide, na.rm = TRUE),

.groups = "drop"

) %>%

mutate(across(where(is.numeric),

~ formatC(.x, digits = 6, format = "f")))

\## Pyrimethanil

Pyrimethanil_descriptive <- quantity %>%

filter(Pyrimethanil > 0)

Pyrimethanil_descriptive %>%

group_by(Experimental_group) %>%

summarise(

no_rows = length(Pyrimethanil),

average = mean(Pyrimethanil, na.rm = TRUE),

stderror = std(Pyrimethanil),

median	= median(Pyrimethanil, na.rm = TRUE),

Q1	= quantile(Pyrimethanil, 0.25, na.rm = TRUE),

Q3	= quantile(Pyrimethanil, 0.75, na.rm = TRUE),

min	= min(Pyrimethanil, na.rm = TRUE),

max	= max(Pyrimethanil, na.rm = TRUE),

.groups = "drop"

) %>%

mutate(across(where(is.numeric),

~ formatC(.x, digits = 6, format = "f")))

\## Quinoxyfen

Quinoxyfen_descriptive <- quantity %>%

filter(Quinoxyfen > 0)

Quinoxyfen_descriptive %>%

group_by(Experimental_group) %>%

summarise(

no_rows = length(Quinoxyfen),

average = mean(Quinoxyfen, na.rm = TRUE),

stderror = std(Quinoxyfen),

median	= median(Quinoxyfen, na.rm = TRUE),

Q1	= quantile(Quinoxyfen, 0.25, na.rm = TRUE),

Q3	= quantile(Quinoxyfen, 0.75, na.rm = TRUE),

min	= min(Quinoxyfen, na.rm = TRUE),

max	= max(Quinoxyfen, na.rm = TRUE),

.groups = "drop"

) %>%

mutate(across(where(is.numeric),

~ formatC(.x, digits = 6, format = "f")))

\## Spinosad-A

SpinosadA_descriptive <- quantity %>%

filter(SpinosadA > 0)

SpinosadA_descriptive %>%

group_by(Experimental_group) %>%

summarise(

no_rows = length(SpinosadA),

average = mean(SpinosadA, na.rm = TRUE),

stderror = std(SpinosadA),

median	= median(SpinosadA, na.rm = TRUE),

Q1	= quantile(SpinosadA, 0.25, na.rm = TRUE),

Q3	= quantile(SpinosadA, 0.75, na.rm = TRUE),

min	= min(SpinosadA, na.rm = TRUE),

max	= max(SpinosadA, na.rm = TRUE),

.groups = "drop"

) %>%

mutate(across(where(is.numeric),

~ formatC(.x, digits = 6, format = "f")))

\## Tebuconazole

Tebuconazole_descriptive <- quantity %>%

filter(Tebuconazole > 0)

Tebuconazole_descriptive %>%

group_by(Experimental_group) %>%

summarise(

no_rows = length(Tebuconazole),

average = mean(Tebuconazole, na.rm = TRUE),

stderror = std(Tebuconazole),

median	= median(Tebuconazole, na.rm = TRUE),

Q1	= quantile(Tebuconazole, 0.25, na.rm = TRUE),

Q3	= quantile(Tebuconazole, 0.75, na.rm = TRUE),

min	= min(Tebuconazole, na.rm = TRUE),

max	= max(Tebuconazole, na.rm = TRUE),

.groups = "drop"

) %>%

mutate(across(where(is.numeric),

~ formatC(.x, digits = 6, format = "f")))

\## Thiacloprid

Thiacloprid_descriptive <- quantity %>%

filter(Thiacloprid > 0)

Thiacloprid_descriptive %>%

group_by(Experimental_group) %>%

summarise(

no_rows = length(Thiacloprid),

average = mean(Thiacloprid, na.rm = TRUE),

stderror = std(Thiacloprid),

median	= median(Thiacloprid, na.rm = TRUE),

Q1	= quantile(Thiacloprid, 0.25, na.rm = TRUE),

Q3	= quantile(Thiacloprid, 0.75, na.rm = TRUE),

min	= min(Thiacloprid, na.rm = TRUE),

max	= max(Thiacloprid, na.rm = TRUE),

.groups = "drop"

) %>%

mutate(across(where(is.numeric),

~ formatC(.x, digits = 6, format = "f")))

\## Thiametoxam

Thiametoxam_descriptive <- quantity %>%

filter(Thiametoxam > 0)

Thiametoxam_descriptive %>%

group_by(Experimental_group) %>%

summarise(

no_rows = length(Thiametoxam),

average = mean(Thiametoxam, na.rm = TRUE),

stderror = std(Thiametoxam),

median	= median(Thiametoxam, na.rm = TRUE),

Q1	= quantile(Thiametoxam, 0.25, na.rm = TRUE),

Q3	= quantile(Thiametoxam, 0.75, na.rm = TRUE),

min	= min(Thiametoxam, na.rm = TRUE),

max	= max(Thiametoxam, na.rm = TRUE),

.groups = "drop"

) %>%

mutate(across(where(is.numeric),

~ formatC(.x, digits = 6, format = "f")))

\## Tolyfluanid

Tolyfluanid_presence <- fisher[,c(3,37)]

Tolyfluanid_quantity <- quantity[,c(3,37)]

chisq.test(table(Tolyfluanid_presence))

Tolyfluanid_detected <- Tolyfluanid_presence %>%

filter(Tolyfluanid > 0)

Tolyfluanid_detected %>%

group_by(Experimental_group) %>%

summarise(no_rows = length(Tolyfluanid))

Tolyfluanid_quantified <- Tolyfluanid_quantity %>%

filter(Tolyfluanid > 0)

wilcox.test(Tolyfluanid ~ Experimental_group, data=Tolyfluanid_quantified)

Tolyfluanid_quantified %>%

group_by(Experimental_group) %>%

summarise(

no_rows = length(Tolyfluanid),

average = mean(Tolyfluanid, na.rm = TRUE),

stderror = std(Tolyfluanid),

median	= median(Tolyfluanid, na.rm = TRUE),

Q1	= quantile(Tolyfluanid, 0.25, na.rm = TRUE),

Q3	= quantile(Tolyfluanid, 0.75, na.rm = TRUE),

min	= min(Tolyfluanid, na.rm = TRUE),

max	= max(Tolyfluanid, na.rm = TRUE),

.groups = "drop"

) %>%

mutate(across(where(is.numeric),

~ formatC(.x, digits = 6, format = "f")))

\## Trifluarin

Trifluarin_descriptive <- quantity %>%

filter(Trifluarin > 0)

Trifluarin_descriptive %>%

group_by(Experimental_group) %>%

summarise(

no_rows = length(Trifluarin),

average = mean(Trifluarin, na.rm = TRUE),

stderror = std(Trifluarin),

median	= median(Trifluarin, na.rm = TRUE),

Q1	= quantile(Trifluarin, 0.25, na.rm = TRUE),

Q3	= quantile(Trifluarin, 0.75, na.rm = TRUE),

min	= min(Trifluarin, na.rm = TRUE),

max	= max(Trifluarin, na.rm = TRUE),

.groups = "drop"

) %>%

mutate(across(where(is.numeric),

~ formatC(.x, digits = 6, format = "f")))

\## Triflusulfuron-methyl

Triflusulfuron_methyl_descriptive <- quantity %>%

filter(Triflusulfuron_methyl > 0)

Triflusulfuron_methyl_descriptive %>%

group_by(Experimental_group) %>%

summarise(

no_rows = length(Triflusulfuron_methyl),

average = mean(Triflusulfuron_methyl, na.rm = TRUE),

stderror = std(Triflusulfuron_methyl),

median	= median(Triflusulfuron_methyl, na.rm = TRUE),

Q1	= quantile(Triflusulfuron_methyl, 0.25, na.rm = TRUE),

Q3	= quantile(Triflusulfuron_methyl, 0.75, na.rm = TRUE),

min	= min(Triflusulfuron_methyl, na.rm = TRUE),

max	= max(Triflusulfuron_methyl, na.rm = TRUE),

.groups = "drop"

) %>%

mutate(across(where(is.numeric),

~ formatC(.x, digits = 6, format = "f")))

#Descriptive statistics (Table 3)

mean(data$nb_compose_sang)

min(data$nb_compose_sang)

max(data$nb_compose_sang)

std(data$nb_compose_sang)

mean(organic$nb_compose_sang)

min(organic$nb_compose_sang)

max(organic$nb_compose_sang)

std(organic$nb_compose_sang)

mean(conventional$nb_compose_sang)

min(conventional$nb_compose_sang)

max(conventional$nb_compose_sang)

std(conventional$nb_compose_sang)

mean(data$sum_pesticides_scale)

min(data$sum_pesticides_scale)

max(data$sum_pesticides_scale)

std(data$sum_pesticides_scale)

mean(organic$sum_pesticides_scale)

min(organic$sum_pesticides_scale)

max(organic$sum_pesticides_scale)

std(organic$sum_pesticides_scale)

mean(conventional$sum_pesticides_scale)

min(conventional$sum_pesticides_scale)

max(conventional$sum_pesticides_scale)

std(conventional$sum_pesticides_scale)

mean(data$TI_tot_indiv)

min(data$TI_tot_indiv)

max(data$TI_tot_indiv)

std(data$TI_tot_indiv)

mean(organic$TI_tot_indiv)

min(organic$TI_tot_indiv)

max(organic$TI_tot_indiv)

std(organic$TI_tot_indiv)

mean(conventional$TI_tot_indiv)

min(conventional$TI_tot_indiv)

max(conventional$TI_tot_indiv)

std(conventional$TI_tot_indiv)

\### AIC selection and model averaging

npp_analyses <- glm(Nppp ~ Sex + Experimental_group + Body_condition + Sex : Experimental_group , data=data, family="poisson", na.action=na.fail)

Anova(npp_analyses, type="3")

summary(npp_analyses)

npp_stand <- standardize(npp_analyses)

npp_dredge <- dredge(npp_analyses, trace=F, rank="AICc")

npp_dredge

a <- get.models(npp_dredge, subset = delta < 2)

summary(model.avg(a, confint = T))

confint(model.avg(a))

sumpestscale_analyses <- lm(sum_pesticides_scale ~ Sex + Experimental_group + Body_condition + Sex : Experimental_group , data=data, na.action=na.fail)

Anova(sumpestscale_analyses, type="3")

summary(sumpestscale_analyses)

sumpestscale_stand <- standardize(sumpestscale_analyses)

sumpestscale_dredge <- dredge(sumpestscale_stand, trace=F, rank="AICc")

sumpestscale_dredge

a <- get.models(sumpestscale_dredge, subset = delta < 2)

summary(model.avg(a, confint = T))

confint(model.avg(a))

titot_analyses <- lm(TI_tot ~ Sex + Experimental_group + Body_condition + Sex : Experimental_group , data=data, na.action=na.fail)

Anova(titot_analyses, type="3")

summary(titot_analyses)

titot_stand <- standardize(titot_analyses)

titot_dredge <- dredge(titot_stand, trace=F, rank="AICc")

titot_dredge

a <- get.models(titot_dredge, subset = delta < 2)

summary(model.avg(a, confint = T))

confint(model.avg(a))

\## Figure 3A,B,C

ggplot(data=data, aes(x=Experimental_group, y=nb_compose_sang, fill=Experimental_group))+

geom_boxplot()+

scale_fill_manual(values = c("tomato3", "darkolivegreen3")) +

theme_bw()+

theme(

panel.border = element_blank(),

panel.grid.major = element_blank(),

panel.grid.minor = element_blank(),

axis.line = element_line(colour = "black"),

axis.text.x= element_text(color="black", size="24"),

axis.text.y= element_text(color="black", size="24"),

axis.title.x = element_text(face="bold", size="26"),

axis.title.y = element_text(color="black", size="26"),

legend.position = "none"

)+

labs(y=expression(Number~of~PPP~(N[PPP])),

x="Experimental group")

ggplot(data=data, aes(x=Experimental_group, y=sum_pesticides_scale, fill=Experimental_group))+

geom_boxplot()+

scale_fill_manual(values = c("tomato3", "darkolivegreen3")) +

theme_bw()+

theme(

panel.border = element_blank(),

panel.grid.major = element_blank(),

panel.grid.minor = element_blank(),

axis.line = element_line(colour = "black"),

axis.text.x= element_text(color="black", size="24"),

axis.text.y= element_text(color="black", size="24"),

axis.title.x = element_text(face="bold", size="26"),

axis.title.y = element_text(color="black", size="26"),

legend.position = "none"

)+

labs(y=expression(Σ*group('[',list(pesticides),']')[scaled]),

x="Experimental group")

ggplot(data=data, aes(x=Experimental_group, y=TI_tot_indiv, fill=Experimental_group))+

geom_boxplot()+

scale_fill_manual(values = c("tomato3", "darkolivegreen3")) +

theme_bw()+

theme(

panel.border = element_blank(),

panel.grid.major = element_blank(),

panel.grid.minor = element_blank(),

axis.line = element_line(colour = "black"),

axis.text.x= element_text(color="black", size="24"),

axis.text.y= element_text(color="black", size="24"),

axis.title.x = element_text(face="bold", size="26"),

axis.title.y = element_text(color="black", size="26")

)+

labs(y=expression(Toxic~unit~(TI[tot])),

x="Experimental group")

\###### Analyses of diversity indexes #####

\## Bray-curtis index

concentration <- data[, c(2,14:46)]

concentration$Partridge_ID <- as.factor(concentration$Partridge_ID)

concentration_matrixnew <- concentration %>%

column_to_rownames("Partridge_ID") %>%

as.matrix()

dist_bc <- vegan::vegdist(concentration_matrixnew, "bray")

\## Betadisper analysis for Bray-Curtis index

dispersion_bc <- betadisper(dist_bc, Experimental_group)

anova(dispersion_bc)

\## PERMANOVA analyses

adonis2(dist_bc ~ Experimental_group, data=data, permutations = 1000)

nmds_result_bc <- metaMDS(dist_bc, k = 2)

nmds_coord_bc <- as.data.frame(scores(nmds_result_bc, display = "sites"))

nmds_coord_bc$Experimental_group <- Group

nrow(nmds_coord_bc)

length(Group)

centroids_bc <- aggregate(cbind(NMDS1, NMDS2) ~ Group, data = nmds_coord_bc, FUN = mean)

\# Figure 4A

ggplot(nmds_coord_bc, aes(x = NMDS1, y = NMDS2, color = Group)) +

geom_point(size = 3, alpha = 0.7) +

stat_ellipse(aes(fill = Group), geom = "polygon", alpha = 0.2, level = 0.95) +

geom_point(data = centroids_bc, aes(x = NMDS1, y = NMDS2), size = 5, shape = 18, stroke = 1.5) +

labs(x = "NMDS1", y = "NMDS2", title = "Bray-Curtis Index - Conventional vs Organic treatment") +

scale_color_manual(values = c("tomato3", "darkolivegreen3")) +

scale_fill_manual(values = c("tomato3","darkolivegreen3")) +

theme(

panel.grid.major = element_blank(),

panel.grid.minor = element_blank(),

panel.background = element_blank(),

panel.border = element_blank(),

axis.line = element_line(color = "grey")

)

\## Jaccard index

dist_jc <- vegan::vegdist(concentration_matrixnew, "bray")

\## Betadisper analysis for Bray-Curtis index

dispersion_jc <- betadisper(dist_jc, Experimental_group)

anova(dispersion_jc)

\## PERMANOVA analyses

adonis2(dist_jc ~ Experimental_group, data=data, permutations = 1000)

nmds_result_jc <- metaMDS(dist_jc, k = 2)

nmds_coord_jc <- as.data.frame(scores(nmds_result_jc, display = "sites"))

nmds_coord_jc$Experimental_group <- Group

nrow(nmds_coord_jc)

length(Group)

centroids_jc <- aggregate(cbind(NMDS1, NMDS2) ~ Group, data = nmds_coord_jc, FUN = mean)

\# Figure 4B

ggplot(nmds_coord_jc, aes(x = NMDS1, y = NMDS2, color = Group)) +

geom_point(size = 3, alpha = 0.7) +

stat_ellipse(aes(fill = Group), geom = "polygon", alpha = 0.2, level = 0.95) +

geom_point(data = centroids_jc, aes(x = NMDS1, y = NMDS2), size = 5, shape = 18, stroke = 1.5) +

labs(x = "NMDS1", y = "NMDS2", title = "Bray-Curtis Index - Conventional vs Organic treatment") +

scale_color_manual(values = c("tomato3", "darkolivegreen3")) +

scale_fill_manual(values = c("tomato3","darkolivegreen3")) +

theme(panel.grid.major = element_blank(), panel.grid.minor = element_blank(), panel.background = element_blank(), panel.border = element_blank(), axis.line = element_line(color = "grey"))
